# The effect of antibiotic selection on collateral effects and evolvability of uropathogenic *Escherichia coli*

**DOI:** 10.1101/2023.11.14.567005

**Authors:** Beth James, Hishikha Reesaul, Sidra Kashif, Mahboobeh Behruznia, Conor J. Meehan, Maria Rosa Domingo-Sananes, Alasdair T. M. Hubbard

## Abstract

Trimethoprim is recommended as a first-line treatment of urinary tract infections (UTIs) in the UK. In 2018, 31.4% of *Escherichia coli* isolated from UTIs in England were trimethoprim resistant, leading to overreliance on other first and second-line antibiotics. Here, we assessed whether prior selection with trimethoprim results in collateral effects to other antibiotics recommended for the treatment of UTIs. As collateral effects, we considered changes in susceptibility, mutation-selection window and population establishment probability. We selected 10 trimethoprim-resistant derivatives from three clinical isolates of uropathogenic *Escherichia coli.* We found that mutations conferring trimethoprim resistance did not have any collateral effects to fosfomycin. In contrast, resistance to trimethoprim resulted in decreased susceptibility (collateral resistance) to nitrofurantoin, below the clinical breakpoint, and narrowed the mutation-selection window thereby reducing the maximum concentration for selection of nitrofurantoin resistance mutations. Our analyses demonstrate that multiple collateral responses should be accounted for when predicting and optimising antibiotic use, limiting future AMR emergence.

## Introduction

Urinary tract infections (UTIs) are the fourth most common infection requiring antibiotic treatment, with over 404 million cases and 236,790 deaths being estimated globally in 2019 ^1^. UTIs are a significant economic burden resulting in more than 6 billion dollars (approximately $18 per person in the US) in direct health care costs globally each year ^2^. Importantly, in Europe, there were 48,700 deaths attributed to antimicrobial resistant UTIs in 2019 ^3^.

The predominant cause of UTIs is *Escherichia coli*, with a prevalence of 75% in uncomplicated UTIs ^4^. UTIs caused by *E. coli* are associated with high levels of recurrence of infection (40%) and approximately 40% of patients with *E. coli* bacteraemia are thought to have a urinary source ^5^. Currently, the recommended first-line antibiotics to treat uncomplicated UTIs in the UK are trimethoprim and nitrofurantoin for 3-5 days ^6^. If no improvement in symptoms after initial treatment, second-line antibiotics such as pivmecillinam or fosfomycin are recommended. However, high prevalence of resistance has been identified in pathogens causing UTIs, as found by the English Surveillance Programme for Antimicrobial Utilisation and Resistance (ESPAUR) ^7^. The report found that in 2018 31.9% of all bacterial isolates from UTI samples were resistant to trimethoprim, while 11.7% were resistant to nitrofurantoin and 7.9% were resistant to fosfomycin ^7^. Antimicrobial susceptibility testing is not normally performed in primary care for UTIs prior to prescribing antibiotics and, with the insufficient development of novel antimicrobials continuing, there is a requirement to develop new strategies to maintain efficacy in current antibiotics and optimise their use.

One approach to maintain antibiotic efficacy and limit AMR development is to apply a group of principles collectively known as selection inversion strategies (SIS) ^8^. These exploit the bacterial response to a second antibiotic when resistance to the first is developed and include collateral susceptibility (CS), synergistic interactions and any fitness costs in the presence of the second antibiotic ^8^.

One SIS is to harness CS networks, where resistance to one antibiotic can confer increased susceptibility to another class of antibiotic. CS networks have been previously investigated in a range of bacterial species against a variety of antimicrobial agents ^9–12^. The result of these studies has demonstrated the potential of this strategy to exploit the advantages of CS, via interventions such as drug cycling ^13^ and combinations of antibiotics ^14^. However, there are limitations to the current research into CS. Many current studies on CS have been carried out on laboratory-derived strains and focus on only describing collateral networks without considering antibiotics used clinically ^9^. These studies have provided vital information on the potential benefits of CS, but do not reflect clinical applications or clinically relevant antibiotics. Furthermore, while CS has been investigated, with CS networks identified, these studies have not addressed or considered if these networks increase or decrease the probability of selecting for AMR against a second antibiotic.

A second collateral effect which could be considered to design SIS is the mutant selection window (MSW) ^15^. This is the antibiotic concentration range in which it is possible to select single-step mutations. The lower boundary of the MSW is often defined as the minimum inhibitory concentration (MIC) which is the lowest concentration of an antibiotic at which cells are unable to grow and at which AMR mutations can be selected. The upper boundary is defined as the mutant prevention concentration (MPC) which is the minimum concentration of an antibiotic that prevents the selection of AMR mutations (or the maximum concentration at which mutations can be selected for). Antibiotic concentrations above the MPC are believed to reduce the potential for development of antimicrobial resistance and have been thoroughly tested through *in vivo* and *in vitro* studies ^12, 16–18^. Furthermore, the MPC and width of the MSW can be regarded as a measure of the capacity of strains to evolve resistance towards an antibiotic (evolvability) ^19^. However, there is still limited research into how prior selection and resistance to one antibiotic affects the MSW of a second antibiotic in the context of clinical antibiotic treatment.

A third, potential collateral effect that could influence SIS is the effect that AMR mutations may have on the ability of bacteria to establish a population ^20^. That is, a single cell from a particular isolate has a specific probability of being able to establish a population. It is therefore possible to determine whether specific AMR mutation(s) will increase or decrease the probability of establishing a population. Moreover, similar to CS, we can assess whether AMR mutation(s) conferring resistance to one antibiotic can affect the population establishment probability (PEP) in the presence of low concentrations of a second antibiotic. As this is a relatively new concept, understanding of the effects of AMR mutations on PE in the presence of antibiotics is limited ^20^.

Interest in understanding collateral effects, such as CS, and changes to MSWs and PEPs, to inform SIS to guide antibiotic regimens that limit the emergence of AMR is increasing. However, most previous studies investigate these collateral responses individually, using laboratory-derived bacterial strains and/or non-clinically relevant antibiotics. Additionally, many studies focus on the effect of individual mutations on collateral responses in a single isolate, rather than any consistent effects across a diverse range of mutations and genetic backgrounds on collateral responses. In this study, we explored collateral effects in 10 *in vitro-*derived trimethoprim-resistant derivatives of three uropathogenic *Escherichia coli* clinical isolates. As collateral effects of AMR mutations, we determine collateral susceptibility/resistance, and shifts in MSW and PEP towards two clinically relevant antibiotics; nitrofurantoin and fosfomycin. Using these data, we aimed to determine whether CS, and MSW and PEP shifts are consistent across the different genetic backgrounds of the clinical strains and whether knowledge of these different effects can be combined to optimise the selection of antibiotics to limit the emergence of AMR.

## Results

### Uropathogenic Escherichia coli clinical isolates

We obtained three clinical isolates of *E. coli* from UTIs: UTI-34, UTI-39 and UTI-59 and sequenced their genomes. UTI-34 and UTI-39 belonged to sequence type (ST) 73, and UTI-59 to ST127. Prediction of AMR genes present in the genome found UTI-34 contained *bla*_TEM-1C_ and *tet(A)*, while UTI-39 was predicted to contain *ant(3’’)-Ia* and *sul1*. In contrast UTI-59 did not contain any predicted AMR genes. Although UTI-34 and UTI-39 were both the same ST, we included both in this study to determine whether the presence of differential horizontally acquired AMR genes would impact consistent collateral responses. Antimicrobial susceptibility testing (AST) identified that all three isolates were susceptible to trimethoprim, nitrofurantoin and fosfomycin according to EUCAST clinical breakpoints (Fig. 1A-C). We then carried out an evolutionary ramp experiment ^21, 22^ (see Supplementary Figure S1) to select four independent trimethoprim-resistant derivatives of the clinical strains, although we were only able to obtain two independent trimethoprim-resistant derivatives of UTI-34 from the four selection experiments. Subsequent rounds of selection for trimethoprim-resistant derivatives of UTI-34 were found not to be derivatives of UTI-34 following whole genome sequencing. Therefore, we obtained 10 independently selected trimethoprim-resistant derivatives for further analysis.

**Figure 1:**
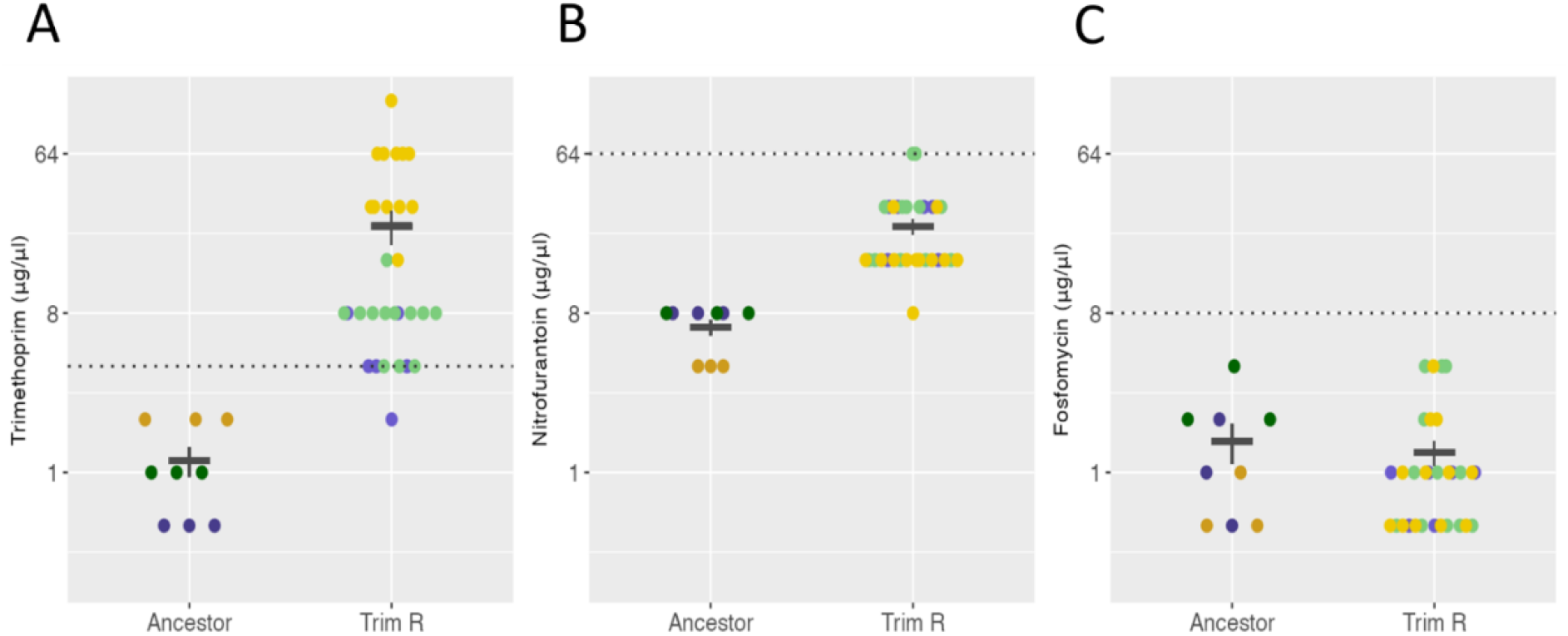
Minimum inhibitory concentrations of ancestor clinical isolates (Ancestor) and trimethoprim-resistant derivatives (Trim-R) for A) trimethoprim, B) nitrofurantoin, and C) fosfomycin. Dotted lines represent the clinical breakpoint for the corresponding antibiotic and error bars represents the standard error of the mean. Colour of data points indicate the ancestor clinical isolate and their linked trimethoprim-resistant derivatives; UTI-34 (blue), UTI-39 (green) and UTI-59 (yellow).

### Phenotypic and genotypic characterisation of trimethoprim-resistant derivatives

Trimethoprim-resistant derivatives of UTI-34 (A-B), UTI-39 (A-D) and UTI-59 (A-D) displayed a significant decrease in susceptibility (p-value = 6.498e-06, Fig. 1A) to trimethoprim following selection (mean MIC = 24.87 µg/ml, min = 2.00 µg/ml, max = 128 µg/ml, Fig. 1A) compared to the ancestors (mean MIC = 1.17 µg/ml, min =0.50 µg/ml, max = 2.00µg/ml Fig. 1A). MICs for all derivatives were at or above the clinical breakpoint of 4 µg/ml and therefore these selected strains are clinically resistant. In addition, we found that overall, the MSW of the trimethoprim-resistant derivatives (MIC-MPC range 24.87 – 144.27 µg/ml, min = 2.00 µg/ml, max 512.00 µg/ml, Fig. 2A) also shifted above the clinical breakpoint compared to the ancestor clinical isolates (P value = 6.516e-06, MIC-MPC range 1.17 – 2.67 µg/ml, min = 0.50 µg/ml, max = 4.00 µg/ml Fig. 2A, see also Figure S4A).

**Figure 2:**
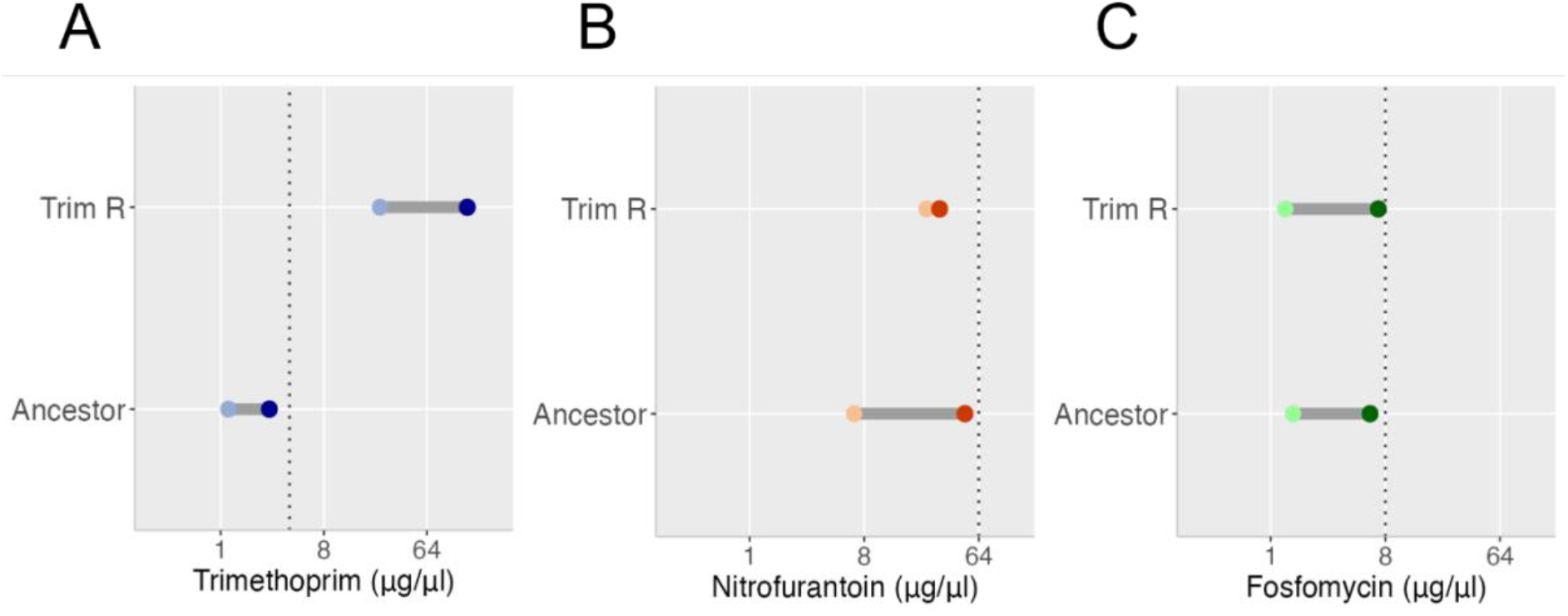
Mutant selection windows of ancestor clinical isolates (Ancestor) and trimethoprim-resistant derivatives (Trim-R) for A) trimethoprim, B) nitrofurantoin, and C) fosfomycin. Dotted lines represent the clinical breakpoint for the corresponding antibiotic. Colour of data points indicate the ancestor clinical isolate and their linked trimethoprim-resistant derivatives, and the lighter dots represent the minimum inhibitory concentration (MIC) and the darker dots represent the mutant prevention concentration (MPC); UTI-34 (blue), UTI-39 (green) and UTI-59 (yellow).

We identified multiple SNPs and/or indels in each of the trimethoprim-resistant derivatives. The majority of SNPs and indels were found in the chromosome, with mutations found in plasmids in 34B (n =3) and 39C (n = 1). Overall, all trimethoprim-resistant derivatives, except 34B, contained 1-2 SNPs or indels in *folA* (the gene encoding the target for trimethoprim) and/or its promoter. Interestingly, 34B had the lowest MIC range (2-4 µg/ml) of the 10 trimethoprim-resistant derivatives. There were identical mutations in *folA* found between the derivatives of a single ancestor clinical isolate, i.e. a Trp30Arg mutation in 39A and 39B, a Leu28Arg and a SNP in the -10 promoter box of *folA* in 59A and 59D and a 4bp insertion in the -10 promoter region of *folA* in 59B and 59C (Fig. 3, Supplementary Table S1). While there is a mutation at the same position in *folA* in 39A and39B (Trp30Arg) and 59C (Trp30Gly), the subsequent amino changes are not identical. Furthermore, there are no identical SNPs and indels between the different ancestor clinical isolate backgrounds. Up to 14 diverse off-target mutations were found in all derivatives with no similarities between the derivatives from different ancestor clinical isolates. Only 59B and 59C carried a single identical off-target mutation in *istB* (Fig. 3, Supplementary Table S1). Other off-target mutations were found in *dnaN*, *dnaA*, *rrf, rppH, rrs, rng*, *ydjL, nfi, pitA*, *garK* and *an* inovirus Gp2 family protein, as well as intragenic regions (Fig. 3, Supplementary Table S1). Importantly, the overall mutational pattern was diverse, with no two derivatives having identical combinations of mutations, demonstrating diversity in the 10 trimethoprim-resistant derivatives (Fig. 3, Supplementary Table S1).

**Figure 3:**
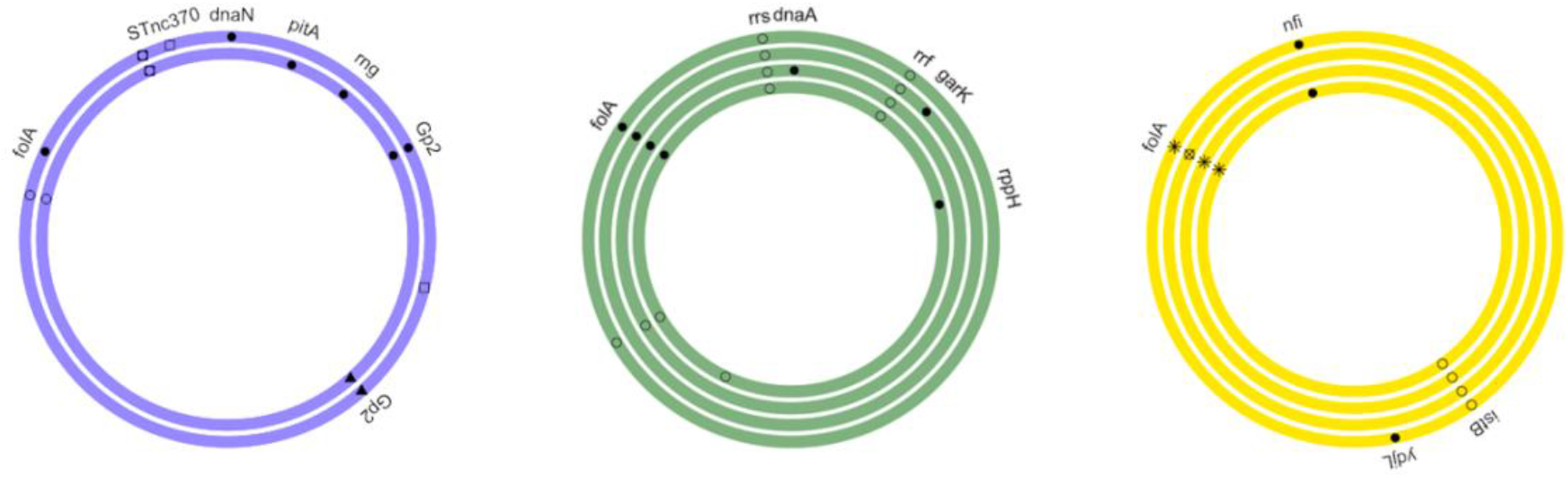
Schematic representing the single nucleotide polymorphisms, insertions and deletions found in the chromosome of trimethoprim-resistant derivatives relative to their individual ancestor clinical isolate. Ring colours indicate the ancestor clinical isolate and their linked trimethoprim-resistant derivatives; UTI-34 (blue), UTI-39 (green) and UTI-59 (yellow). Filled circles represent coding, non-synonymous mutations, open circles represent coding, synonymous or intragenic mutations, filled triangle represent multiple synonymous or non-synonymous mutations, open square represents intragenic indels, asterisk represent *folA* coding and promoter region mutations and circle with cross represents *folA* promoter region mutations only.

Finally, we assessed the effect of the above mutations on fitness of the trimethoprim-derivatives relative to the ancestor clinical isolates (Supplementary Figure S2). We found that there was no significant difference in the maximum growth rate (*r*) of the trimethoprim-resistant derivatives (mean = 0.9987, P value = 0.8338, Fig. 4). However, we identified a significant decrease in relative carrying capacity (*k*, mean = 0.9306, P value = 2.566e-07 Fig. 4) and relative area under the curve (AUC, mean = 0.9028, P value = 6.44e-10, Fig. 4).

**Figure 4:**
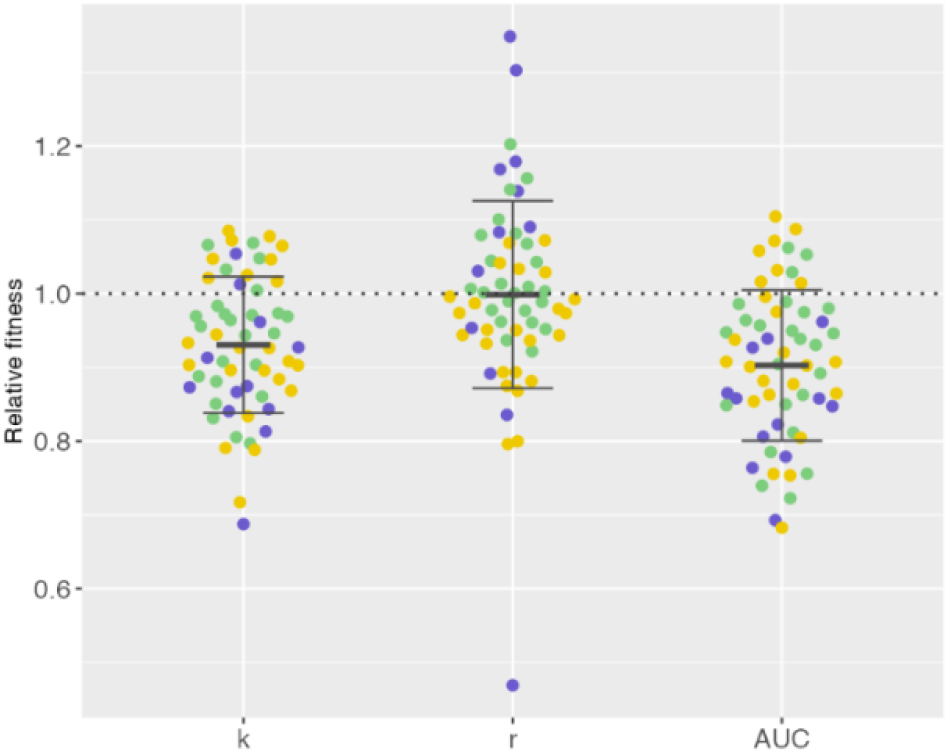
Plots of maximum growth rate (*r*), relative carrying capacity (*k*) and relative area under the curve (AUC) of the trimethoprim-resistance derivatives relative to the ancestor clinical isolates. Error bars represents standard deviation and colour of data points indicate the ancestor clinical isolate and their linked trimethoprim-resistant derivatives; UTI-34 (blue), UTI-39 (green) and UTI-59 (yellow).

### Collateral susceptibility and resistance

Next, we assessed any collateral effects of mutations identified in the trimethoprim-resistant derivatives conferred on the activity of an alternative first-line antibiotic, nitrofurantoin, and a second-line antibiotic, fosfomycin. While strain/derivative level differences were observed (Supplementary Figure S3), since we had a diverse range of mutational profiles in three different genetic backgrounds, we assessed whether there was any consistency in the effects of trimethoprim resistance on CS. We found no significant difference (P value 0.4935) between the MICs of the ancestor clinical isolates (mean MIC = 1.50 µg/ml, min = 0.50 µg/ml, max = 4.00 µg/ml, Fig. 1C) and the trimethoprim-resistant derivatives (mean MIC = 1.30 µg/ml, min = 0.50 µg/ml, max = 4.00 µg/ml Fig. 1C) for fosfomycin; therefore, all were below the clinical breakpoint (8 µg/ml). In contrast, the trimethoprim resistant derivatives displayed significant (P value = 6.498e-06) decreased susceptibility to nitrofurantoin (mean MIC = 24.80 µg/ml, min = 8.00 µg/ml, max = 64 µg/ml Fig. 1B) compared to the ancestor clinical isolates (mean MIC = 6.67 µg/ml, min = 4.00 µg/ml, max = 8.00 µg/ml Fig. 1B). Despite this significant decrease in susceptibility, all MICs for the trimethoprim-resistant derivatives were at or below the clinical breakpoint for nitrofurantoin of 64 µg/ml.

### Mutant selection windows

In this study we used a modified minimum bactericidal concentration assay to assess the MPC for all isolates. While other MPC assays have been established and widely used to determine MSW, we wanted to ensure standardisation of the initial bacterial population in each assay due to the significant decrease in carrying capacity observed in the trimethoprim-resistant derivatives. Additionally, we wanted to assess the MPC in relation to clinical breakpoints set by EUCAST. Again, while strain and derivative level differences were observed in the MSW (Supplementary Figure S4), we assessed any consistent effects. We found that the MSW of trimethoprim-resistant derivatives (MIC-MPC range 1.30 – 7.03 µg/ml, min = 0.50 µg/ml, max = 64.00 µg/ml, Fig. 2C) was not significantly wider than that of the ancestor clinical isolates (P value = 0.9714, MIC-MPC range 1.50 – 6.06 µg/ml, min = 0.5 µg/ml, max = 16.00 µg/ml Fig. 2C) for fosfomycin. In contrast, we found that the MSW for trimethoprim-resistant derivatives (MIC-MPC range 24.80 – 31.47 µg/ml, min = 8.00 µg/ml, max = 64 µg/ml Fig. 2B) was significantly narrower than the ancestor clinical isolates (P value = 8.485e-06, MIC-MPC range 6.67 – 49.78 µg/ml, min = 4 µg/ml, max = 64µg/ml Fig. 2B) for nitrofurantoin. This suggests that prior trimethoprim resistance reduces the window for selection of resistance to nitrofurantoin while does not significantly change the window for selection to fosfomycin.

### Population establishment

As previously observed with both CS and MSWs, we identified strain and derivative level differences in the ability of the trimethoprim-resistant derivatives to be able to establish a population from 1-2 cells relative to the ancestor clinical isolate (Supplementary Figure S5). When considering any consistent effects, we found that while the ability to establish a population decreased in the trimethoprim-resistant derivatives compared to the ancestor (nitrofurantoin mean = 26.8 and fosfomycin mean = 21.0) in the presence of 1/16^th^ MIC of nitrofurantoin (mean = 25.2, Kruskal-Wallis chi-squared = 0.18837, df = 1, P value = 0.6643, Fig. 5A) and fosfomycin (mean = 20.8, Kruskal-Wallis chi-squared = 0.0044544, df = 1, P value = 0.9468, Fig. 5B), this was not statistically significant.

**Figure 5:**
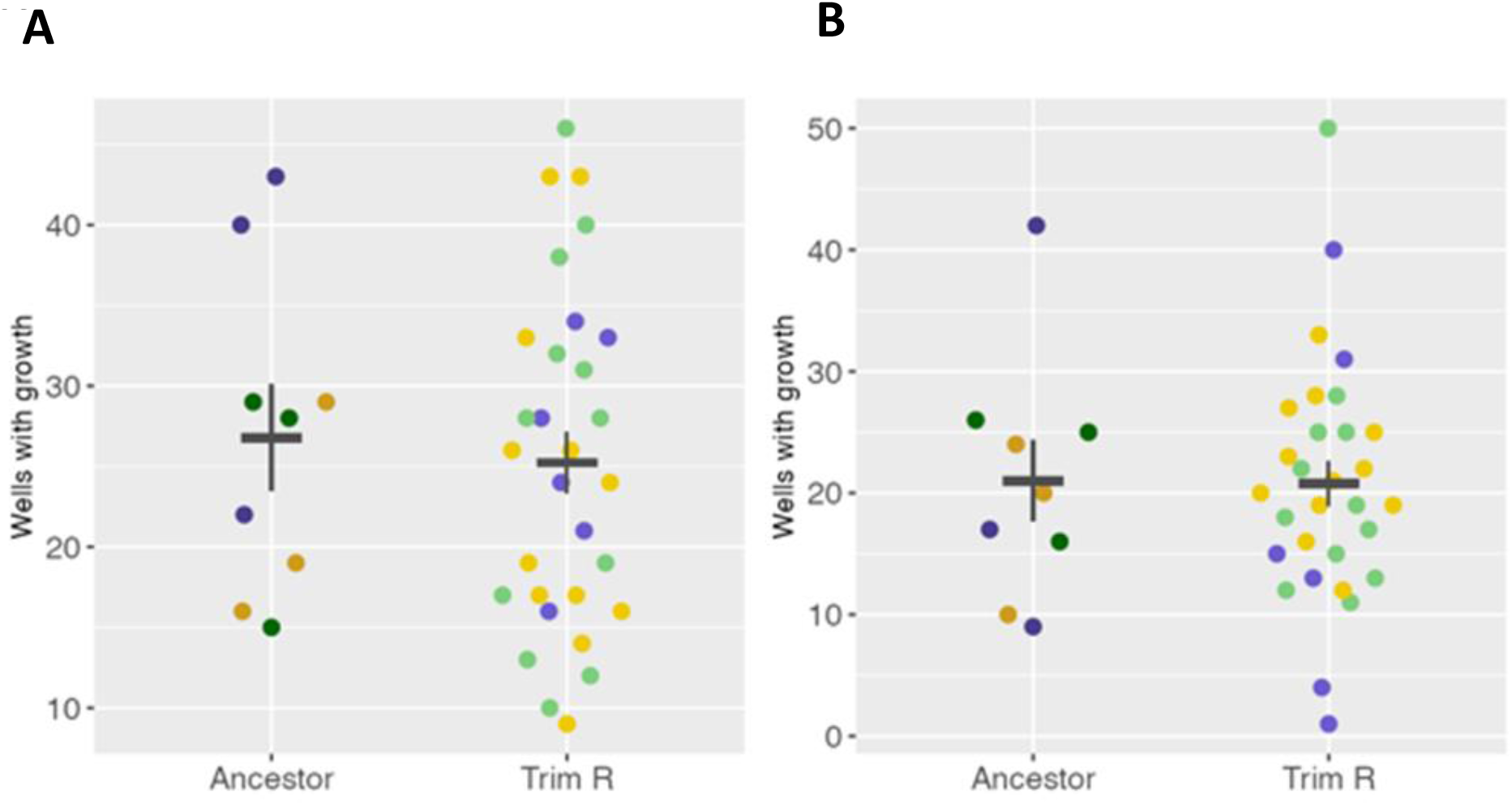
Population establishment of the ancestor clinical isolates and the trimethoprim-resistance derivatives in the presence of either A) nitrofurantoin or B) fosfomycin. Error bars represents standard error of the mean and colour of data points indicate the ancestor clinical isolate and their linked trimethoprim-resistant derivatives; UTI-34 (blue), UTI-39 (green) and UTI-59 (yellow).

## Discussion

To determine whether collateral responses could be used to inform antimicrobial prescribing, it is important to assess this in clinical isolates with resistance to a first-line antibiotic to which resistance is common. Trimethoprim is a common first-line antibiotic recommended for the treatment of uncomplicated UTIs in non-pregnant women 16 years old and over in the UK ^6^. The incidence of trimethoprim resistance in UTIs is high, with 31.9% of all bacterial isolates and 31.4% of *E. coli* from UTI samples resistant to trimethoprim in England in 2018 ^7^. In this study, we selected for 10 trimethoprim-resistant derivatives from three uropathogenic *E. coli* strains to determine possible collateral effects, namely CS, MSWs and PEP, to nitrofurantoin and fosfomycin. All three antibiotics are recommended first- or second-line antibiotics for the treatment of uncomplicated UTIs in the UK.

We used a 72-hour evolutionary ramp experiment to select for resistance to closely match the recommended 3-day course of treatment for uncomplicated UTIs with trimethoprim ^6^. We found the trimethoprim-resistant derivatives selected for *in vitro* had MICs which were at or above the clinical breakpoint for trimethoprim, confirming selection of clinical resistance. We also observed a shift in the MSW for trimethoprim above the clinical breakpoint, suggesting that continued exposure to trimethoprim may increase the probability of selection of higher-level resistance to the antibiotic. It is also possible that derivatives with higher levels of resistance were not observed in our evolutionary ramp due to high fitness costs of mutations conferring high level resistance in this competitive environment. SNPs and indels were identified in *folA* or its promoter ^23^ were found in all but one of the trimethoprim-resistant derivatives. This gene produces dihydrofolate reductase, which synthesises the co-enzyme tetrahydrofolic acid, and the primary target of trimethoprim ^24^. Many of these mutations have previously been identified in following *in vitro* selection of trimethoprim resistance and likely either alter the affinity of the antibiotic to the dihydrofolate reductase or results in the hyperproduction of the enzyme ^12^. Finally, despite the diverse range of *folA* and off-target mutations present within the 10 trimethoprim-resistant derivatives, we did not observe any consistent effect on maximum growth rate compared to the ancestor clinical isolates. While specific mutations and mutational profiles often have varying effects on fitness, it is expected that overall, these would have a deleterious effect on fitness ^25^. Consistent with this, although we did not observe decreased growth rates, we did observe an overall decrease in the relative carrying capacity of the trimethoprim-resistant derivatives. Additionally, the unique mutational profile of each of the trimethoprim-resistant derivatives, especially the diverse off-target mutations, may contribute to the derivative level variation in CS, MSW and PE.

When applying consistent collateral effects to inform SIS such as antibiotic cycling, it is important to consider antibiotics which are recommended alternatives for the treatment of, in this case, UTIs. Nitrofurantoin is recommended as an alternative first-line treatment for uncomplicated UTIs in non-pregnant women 16 years old and over in the UK, while fosfomycin is a recommended second-line treatment for the same patient group ^6^. Fosfomycin is a broad-spectrum antibiotic that targets mucopeptide synthesis, a part of the synthesis of peptidoglycan, a key precursor in the formation of bacterial cell walls ^26^. Of the SNPs and indels identified within the trimethoprim-resistant derivatives none were known to produce resistance to fosfomycin. We did not observe any significant consistent collateral responses to CS, MSW and PEP in the trimethoprim-resistant derivatives to fosfomycin. This is consistent with other studies which have found there to be minimal cross-resistance between fosfomycin and other antibiotics, due to the unique structure of fosfomycin ^27^. Due to lack of changes in the concentration range required to select for resistant mutations and ability to establish a population, as well as lack of collateral resistance, fosfomycin is likely to be a good alternative antibiotic to use if mutational resistance to trimethoprim is observed.

Consistent collateral resistance in nitrofurantoin-resistant *E. coli* towards ciprofloxacin, amoxicillin and azithromycin have previously been documented ^12^. However, consistent collateral responses between trimethoprim-resistant *E. coli* and nitrofurantoin have not yet been studied. We found that while collateral resistance was identified in the trimethoprim-resistant derivatives towards nitrofurantoin, the MPC was lower relative to the ancestor clinical isolates, narrowing the MSW. This suggests that relative to the ancestor clinical isolates, there is a reduced concentration window for the selection of mutations conferring nitrofurantoin resistance.

As previously seen with fosfomycin, there is no significant difference in the ability for the trimethoprim-resistant derivatives to establish a population in the presence of nitrofurantoin compared to the ancestor clinical isolates. Collateral resistance is usually a response to avoid, but as it is below the clinical breakpoint, the trimethoprim-resistant derivatives are therefore clinically susceptible. This, coupled with a narrower MSW, indicates that nitrofurantoin may still be a good alternative antibiotic to use if mutational resistance to trimethoprim is observed. Although, in contrast to fosfomycin, mutational resistance to trimethoprim may reduce the probability for future selection of nitrofurantoin resistance, thereby limiting the development of further AMR. Nitrofurantoin has a diverse mode of action, following cytochrome P450 reductase mediated activation of the nitro group, it is known to inhibit the activity of ribosomes, protein synthesis machinery, various enzymes involved in the synthesis of carbohydrates, cell wall and DNA ^28^. While several of the mutations in genes involved in protein synthesis and DNA replication and repair were identified in the trimethoprim-resistant derivatives, they have not previously been linked to decreased susceptibility to nitrofurantoin. Therefore, the mechanism of collateral resistance is unclear but could be due to some combination of the off-target mutations observed.

We acknowledge that this study has used a limited number of clinical isolates to obtain trimethoprim-resistant derivatives from, two of which belong to the same sequence type. However, as UTI-34 and UTI-39 have different AMR gene profiles, it is important to assess the effects of different combination of AMR genes on collateral responses. In addition, the mutational profile of all 10 trimethoprim-derivatives were diverse and therefore represents a good range of isolates to assess collateral responses. Also, all experiments were performed in laboratory growth media. While this important for standardisation across laboratories, it is not representative of a physiological environment and evidence is mounting which demonstrates the environment is important for assessing AMR ^29^. This is especially important when considering nitrofurantoin activity is enhanced in acidic conditions ^30^. It will therefore be important to study collateral responses in physiologically relevant environments and with a diverse range of isolates to confirm observations before they can be used to inform SIS and clinical interventions.

## Conclusions

This study has demonstrated consistent collateral responses which arise from pre-selected trimethoprim resistance in uropathogenic *E. coli.* Specifically, we have shown the pre-selected resistance to trimethoprim lead to no collateral responses to fosfomycin while causing collateral resistance and a narrower MSW to nitrofurantoin. This suggests that identifying consistent collateral responses may be used to design antibiotic regimens which improve treatment options and outcomes, as well as reduce the development of AMR. However, these collateral responses need to be assessed together and not in isolation to produce a full picture of potential consistent collateral responses.

## Materials and methods

### Bacterial strains, growth conditions and antibiotics

Three clinical isolates of *E. coli,* isolated from urine of patients with UTIs, were obtained from Nottingham University Hospitals NHS Trust Pathogen Bank. Strains 19Y000034 (UTI-34), 19Y000039 (UTI-39) and 19Y000059 (UTI-59) were selected as antimicrobial susceptibility testing via disc diffusion indicated they were susceptible to all recommended antibiotics for treatment of UTIs.

Unless otherwise stated, all bacterial strains were plated on Muller Hinton (MH) agar (Millipore, USA) and incubated at 37°C for 18 hours. Following this, starter cultures were made from a single bacterial colony in 10 ml of MH broth and incubated at 37°C, 150 rpm for 18 hours.

All the following antibiotics and chemicals were solubilised to a stock concentration of 10 mg/ml and sterile filtered through a 0.22 µM polyethersulfone filter unit (Millipore, USA).

Trimethoprim and nitrofurantoin (Sigma, USA) were both solubilised in dimethyl sulfoxide, (DMSO, Sigma, USA). Fosfomycin (Cambridge Bioscience, UK) was solubilised in deionised water and sonicated in an ultrasonic water bath (Branson Ultrasonics Corporation, USA) for 30 minutes. Glucose-6-phosphate (Sigma, USA) was solubilised in deionised water.

### Antimicrobial susceptibility testing

Antimicrobial susceptibility was assessed using the broth microdilution assay following EUCAST guidelines (https://www.eucast.org/). All AST assays were performed in MH broth, except fosfomycin where MH broth was supplemented with 25 µg/ml glucose-6-phosphate and incubated at 37°C for 18 hours. Three biological replicates for each strain were produced for each isolate.

### Selection for trimethoprim-resistant mutants

Trimethoprim-resistant derivatives from three clinical isolates of *E. coli* using the evolutionary ramp method ^21, 22^ over 72 hours, to mimic the course of antibiotic treatment (Supplementary Figure S1). Starter cultures were diluted to 1/1000 in both MH broth and MH broth containing ¼ minimum inhibitory concentrations (MIC) of trimethoprim and incubated at 37°C, 150 rpm for 24 hours. The culture with observable growth containing the highest concentration of trimethoprim (e.g. ¼ MIC) was diluted to 1/1000 in MH broth plus the same concentration of trimethoprim (e.g. ¼ MIC) and MH broth with double the concentration of trimethoprim (e.g. ½ MIC) and incubated at 37°C, 150 rpm for 24 hours. This was repeated, with the same concentration of trimethoprim (e.g. ½ MIC) and double the concentration of trimethoprim (e.g. 1 MIC). Following incubation, 100μl of the culture with growth at the highest trimethoprim concentration were plated out on MH agar containing 1x and 2x MIC of trimethoprim and incubated for 18 hours at 37°C. A single colony from MH agar containing 2x MIC of trimethoprim was used to inoculate 10 ml MH broth and incubated at 37°C, 150 rpm for 18 hours. From this, 500μl of this culture was added to 500μl of 20% glycerol and stored at -80°C freezer. For each clinical isolate, four evolutionary ramp experiments were performed to produce four independent lineages.

### Mutant prevention concentration assays

Broth microdilution assays were initially set up as described above for antimicrobial susceptibility testing. From all wells with no visible growth (100 µl total volume per well), 50 µl was plated out onto MH agar (nitrofurantoin and trimethoprim only) and MH agar containing the same corresponding concentration of the appropriate antibiotic. Agar containing fosfomycin was supplemented with 25 µg/ml glucose-6-phosphate. All agar plates were then incubated at 37°C for 18 hours. The mutant prevention concentration (MPC) was determined as the lowest antibiotic concentration containing no visible growth on MH agar plus antibiotics. If growth skipped concentrations (e.g. no growth at 1 µg/ml, growth at 2 µg/ml and no growth at 4 µg/ml) we applied the same interpretation as EUCAST guidelines for broth microdilution assay. Specifically, if one concentration is skipped then the next concentration with no visible growth was recorded as the MPC. However, if more than one concentration is skipped, this is disregarded and MPC was recorded as the lowest concentration with no visible growth. A total of three biological replicates were completed for each isolate.

### Fitness assays

Starter cultures were diluted to an optical density at a wavelength of 600 nm (OD_600_) of 0.1 and furthered diluted 1/1000 in MH broth. Two technical replicates of 100μl of each diluted culture was added to a flat bottom 96-well plate, with two wells containing 100μl of MH broth as negative controls. OD_600_ of each well was read every 10 minutes, with 100 flashes per well in a SPECTROstar NANO microplate reader (BMG Labtech, Germany), incubated at 37°C, 200 rpm for 24 hours. A total of three biological replicates were completed for each isolate. Data was analysed using the GrowthCurver (v0.3.1) ^31^ package in R (v4.3.1) ^32^.

### Probability of population establishment assay

Population establishment assays ^20^ were performed on all isolates in the presence of either nitrofurantoin or fosfomycin. From a starter culture, 100μl diluted in 10ml of MH broth and incubated at 37°C for 2-3 hours, at 150 rpm until they reach an OD_600_ of 0.3-0.6. The cultures were further diluted to an OD_600_ of between 0.1-0.11 in MH broth. These diluted cultures were serially diluted, firstly by 1/1000 in MH broth, and then 1/8000 in MHB plus 1/16 of the MIC of the ancestral isolate of either nitrofurantoin or fosfomycin supplemented with 25 µg/ml glucose-6-phosphate. Following this, 100µl of each culture was added to 57 wells of a 96 well plate individually. Negative controls of 2 x 100µl of MH broth 1/16 of the MIC of the ancestral isolate of either nitrofurantoin or fosfomycin supplemented with 25 µg/ml glucose-6-phosphate were included in each assay, as well as 1 x 100µl of MH broth. Each assay was incubated at 37°C for 24 hours. Finally, each plate was placed in the SPECTROstar NANO or Cytation3 (BioTek, USA) microplate reader at OD_600_ and scanned with 100 flashes. Growth was considered to have occurred in wells where OD_600 >0.1_. Three biological replicates were performed for each isolate-antibiotic combination.

### DNA extractions

DNA was extracted from starter cultures of the three original ancestor isolates and 10 trimethoprim-resistant derivatives using New England Biolabs Monarch Genomic DNA Extraction Kit (New England Biolabs, USA) following the manufacturer’s instructions. All DNA was eluted in molecular grade water and used for long- and short-read sequencing. UTI-59 DNA for long-read sequencing only was extracted using the Wizard HMW DNA Extraction Kit (Promega, USA) following the manufacturer’s instructions excepted eluted in molecular grade water. DNA concentration was quantified using a Qubit fluorometer and 1X dsDNA High Sensitivity (HS) assay kit (ThermoFisher Scientific, USA).

### Whole genome sequencing

Illumina short-read sequencing was provided by MicrobesNG (https://www.microbesng.com). Nanopore sequencing was performed using the native barcoding kit 24 V14 (SQK-NBD114.24 according to the manufacturer’s instruction on the MinION Mk1C platform with R10.4.1 flowcells.

### Bioinformatic analysis

Adapter sequences from the ONT long-read sequences were removed using Porechop (v0.2.4, https://github.com/rrwick/Porechop) with “--end_threshold 95” and “--middle_threshold 85” options, and quality filtered using Filtlong (v0.2.1, https://github.com/rrwick/Filtlong) with default options and using Illumina reads as external reference . Hybrid assembly of the long- and short-read sequences of three ancestor clinical isolates was performed using Unicycler ^33^ (v0.5.0) and annotated with bakta ^34^ (v1.8.1, database v5.0). Identification of single nucleotide polymorphisms (SNPs) within the trimethoprim resistant derivatives were identified using Snippy (v4.6.0, https://github.com/tseemann/snippy). AMR genes predicted to be present in each of the three ancestor clinical isolates were identified using ABRicate (v1.0.1, https://github.com/tseemann/abricate) using the Resfinder ^35^ database. This publication made use of the PubMLST website (https://pubmlst.org/) developed by Keith Jolley and sited at the University of Oxford. The development of that website was funded by the Wellcome Trust. Multi-locus sequence types (MLST) of each of the clinical isolates were performed using MSLT (v2.23.0, https://github.com/tseemann/mlst) using the “ecoli_achtman_4” scheme.

### Statistical analysis

Figures and data analysis were carried out using R ^32^ and Rstudio ^36^. Test Kruskal-Wallis rank sum test was used to test for differences between MIC distributions (approximately log-normal distributed data with ties and unequal sample sizes) and PEP assay results (number of wells with growth, data with ties and unequal sample sizes) For the relative fitness data, Shapiro-Wilk tests were carried out to test for normality. For normally distributed data (k, AUC), a t-test was used to test for deviation from 1. For non-normally distributed data (r), the Wilcoxon signed rank test with continuity correction was used.

## Supporting information

Code and commands

## Data availability

All sequencing reads generated during this project are available in the sequence read archive (SRA) under the BioProject number PRJNA1037559. Specific accession numbers for each isolate are detailed in Supplementary Table S2. All commands and code for bioinformatics analysis and generation of figures, including the datasets for each figure, are available in “Commands_and_Figure_Data.zip”.

## Contributions

The study was conceived by C.J.M., M.D.R.S. and A.T.M.H. The methodology was devised by B.J., M.B., M.D.R.S. and A.T.M.H. Data was collected by B.J., H.R., S.K., M.B. and A.T.M.H. Data analysis was performed by B.J., M.B., M.D.R.S. and A.T.M.H. The original manuscript draft was prepared by B.J. and A.T.M.H and then reviewed and edited by all authors.

