## Supplementary files for "The effect of antibiotic selection on collateral effects and evolvability of uropathogenic *Escherichia coli*"

**Supplementary Table S2:** Accession numbers for each isolate sequenced using the Oxford Nanopore Technologies and/or Illumina sequencing platforms under the BioProject number PRJNA1037559.

| Accession number | Sample Name |
|---|---|
| SAMN38186228 | UTI-34 |
| SAMN38186229 | UTI-39 |
| SAMN38186230 | UTI-59 |
| SAMN38186231 | 34A |
| SAMN38186232 | 34B |
| SAMN38186233 | 39A |
| SAMN38186234 | 39B |
| SAMN38186235 | 39C |
| SAMN38186236 | 39D |
| SAMN38186237 | 59A |
| SAMN38186238 | 59B |
| SAMN38186239 | 59C |
| SAMN38186240 | 59D |

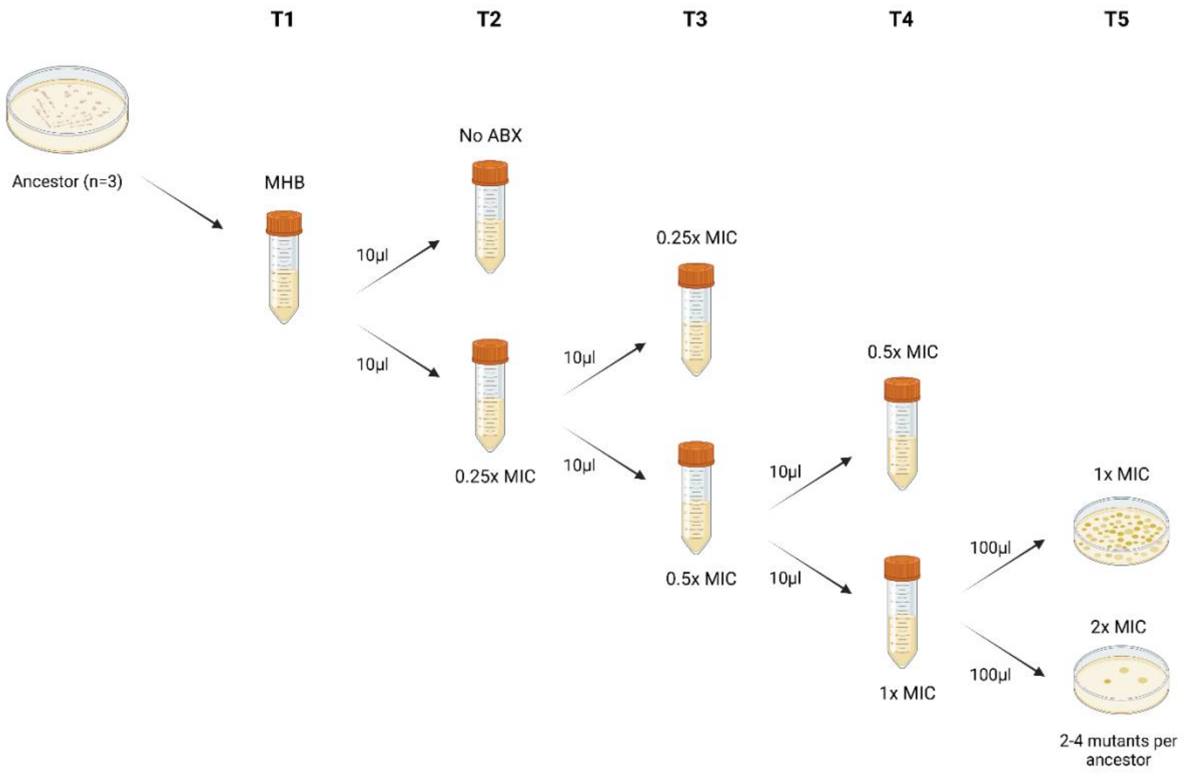


**Supplementary Figure S1:** Diagram of the evolutionary ramp experiment to select for trimethoprim-resistant derivatives from the three clinical isolates of *Escherichia coli*.

A

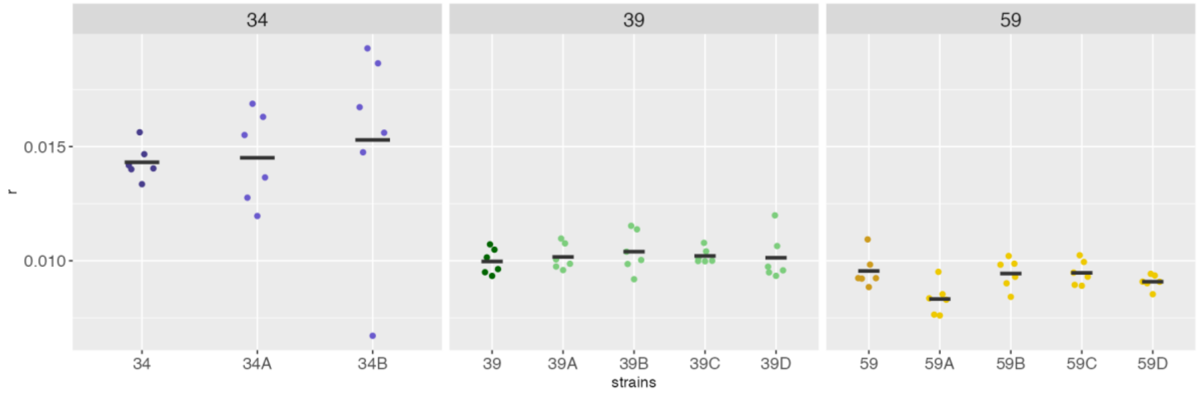


B

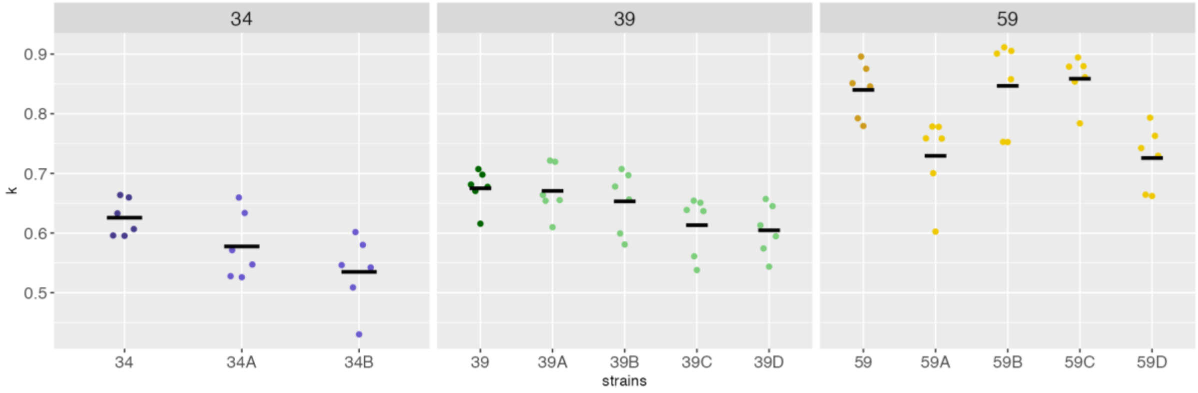


C

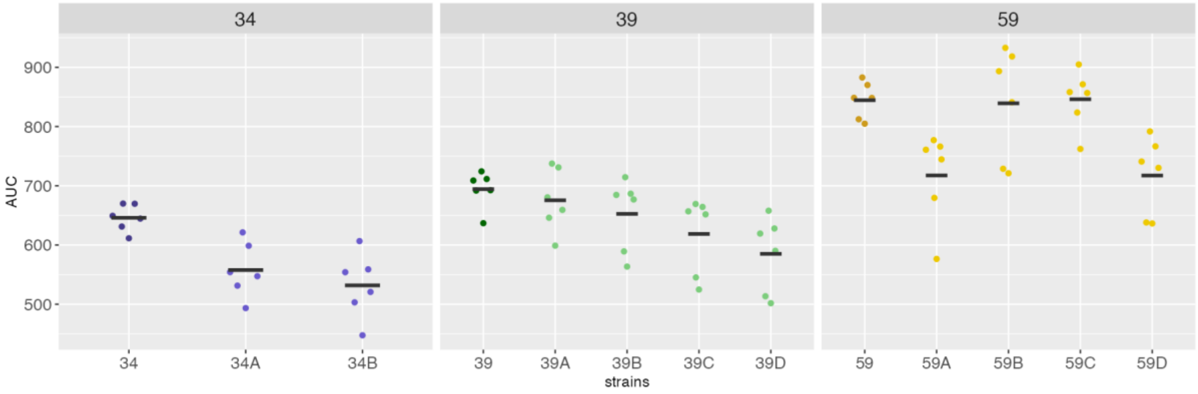


**Supplementary Figure S2:** Plots of A) maximum growth rate (*r*), B) carrying capacity (*k*) and C) area under the curve (AUC) of the trimethoprim-resistance derivatives relative and the ancestor clinical isolates.

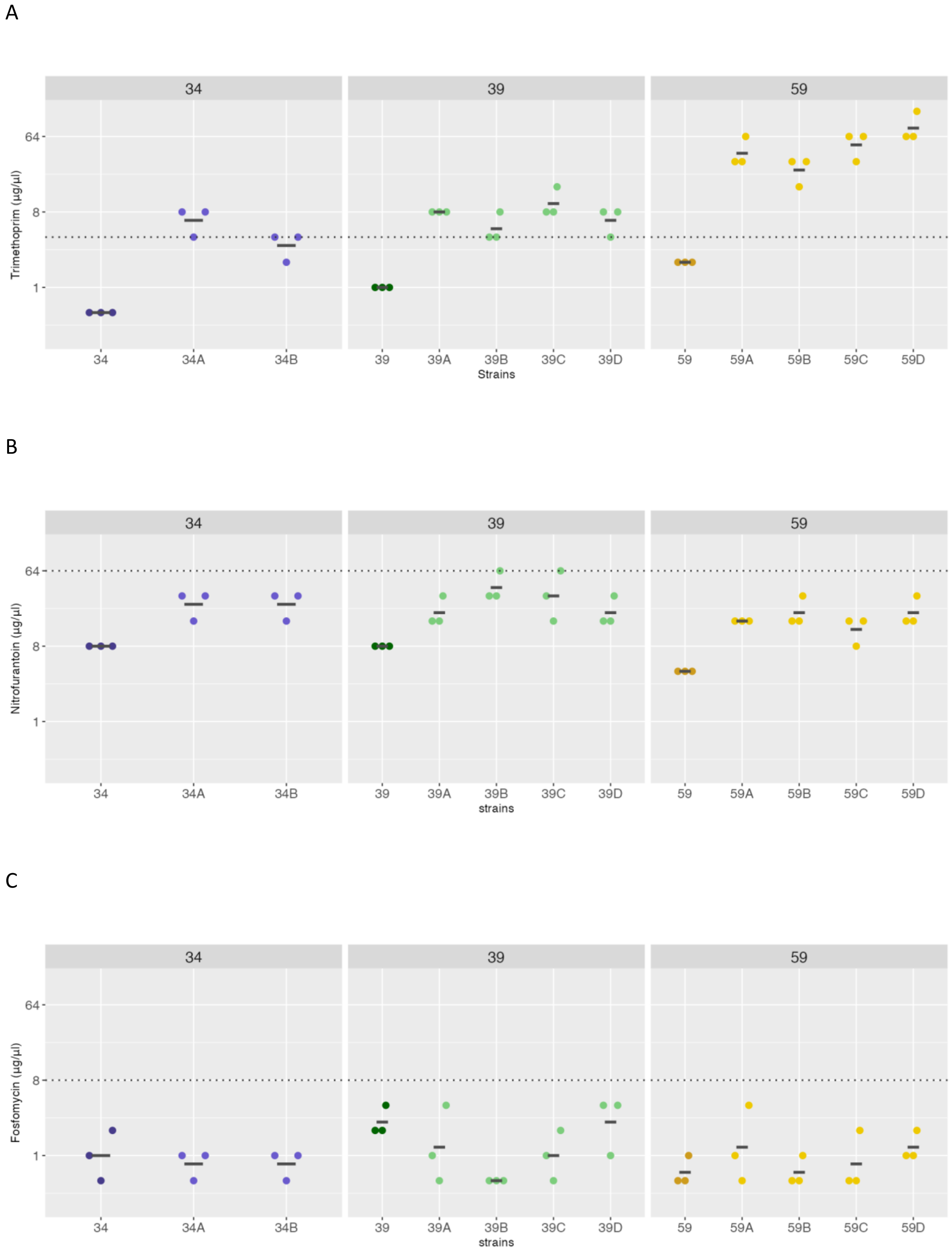


**Supplementary Figure S3:** Minimum inhibitory concentrations of ancestor clinical isolates and trimethoprim-resistant derivatives for A) trimethoprim, B) nitrofurantoin, and C) fosfomycin. Dotted lines represent the clinical breakpoint for the corresponding antibiotic and line for each group represents the mean.

A

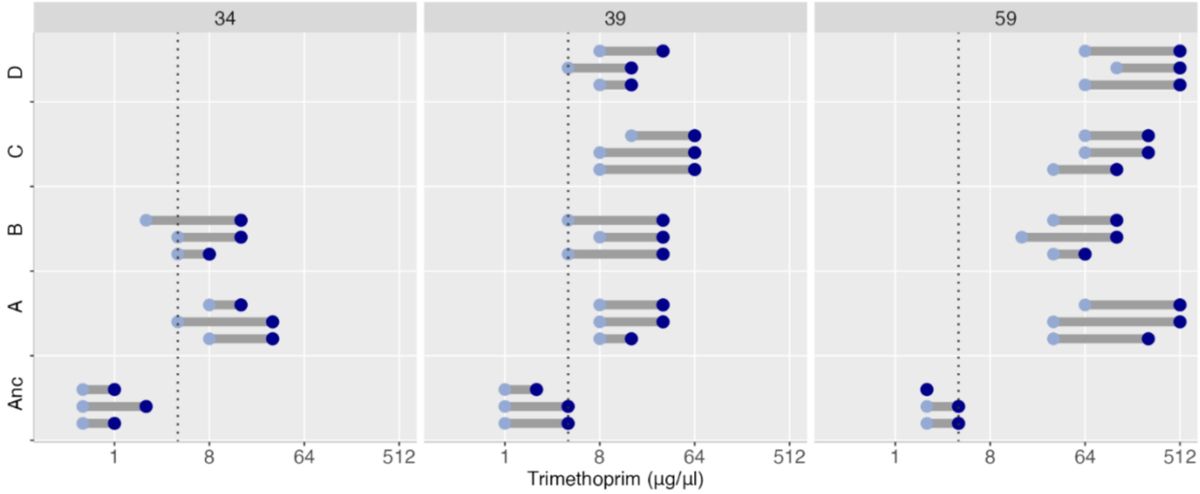


B

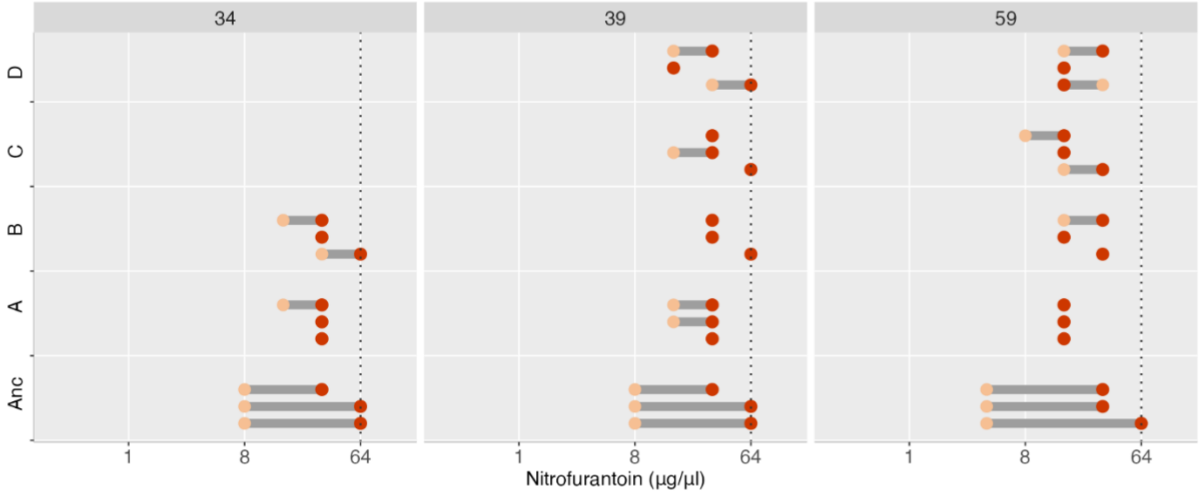


C

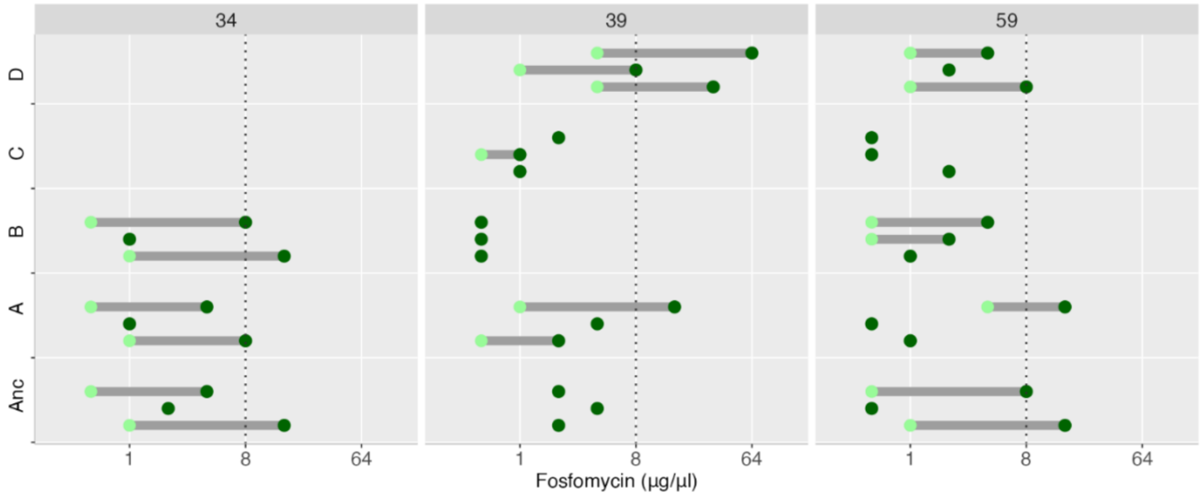


**Supplementary Figure S4:** Mutant selection windows of ancestor clinical isolates (Anc) and trimethoprim-resistant derivatives (A-D) for A) trimethoprim, B) nitrofurantoin, and C) fosfomycin. Dotted lines represent the clinical breakpoint for the corresponding antibiotic. Lighter dots represent the minimum inhibitory concentration (MIC) and the darker dots represent the mutant prevention concentration (MPC).

A

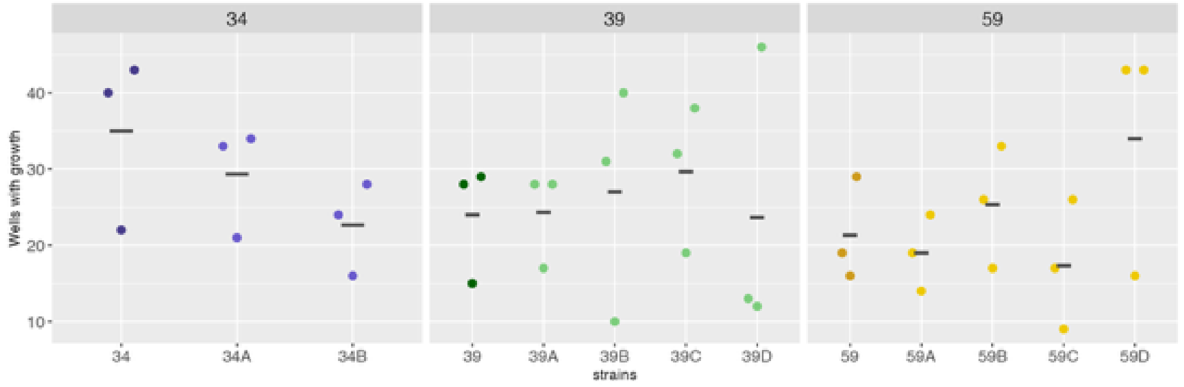


B

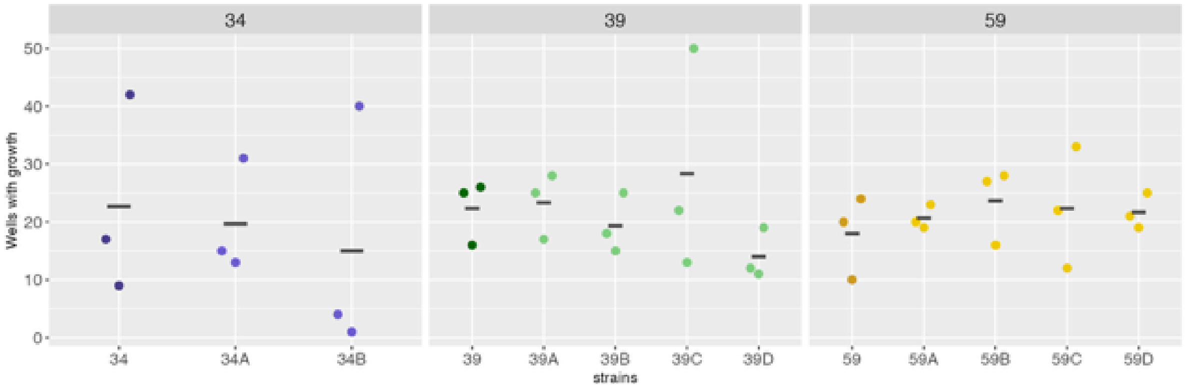


**Supplementary Figure S5:** Population establishment of the trimethoprim-resistance derivatives and the ancestor clinical isolates in the presence of A) nitrofurantoin and B) fosfomycin.
