## Supplementary material for "The effect of antibiotic selection on collateral effects and evolvability of uropathogenic *Escherichia coli*": Code and commands: CirclePlots.R

```
rm(list=ls(all=TRUE))
dev.off()
library(circlize)
setwd('~/') #add path to data files

#genome sizes:
g34 <- 4907664
g39 <- 5137808
g59 <- 5056252

#Plots for strain 34:
#read variant files:
v34a <- read.delim('SNP_data/34A.tsv')
v34a <- v34a[which(v34a$CHROM == "contig_1"),]
v34b <- read.delim('SNP_data/34B.tsv')
v34b <- v34b[which(v34b$CHROM == "contig_1"),]

png("Figures/circle34.png", width = 1500, height = 1500, units = "px", bg = "white",  res = 300)
circos.clear()
circos.par("track.height" = 0.05)
col_text <- "white"
circos.par(start.degree = 90)
circos.initialize(sectors = 'chr', xlim = c(0,g34))
circos.track(ylim=c(0, 1), panel.fun=function(x, y) {
  chr=CELL_META$sector.index
  xlim=CELL_META$xlim
  ylim=CELL_META$ylim
  #circos.axis(labels.cex = 0.6)
}, bg.col="white", bg.border=F, track.height=0.1)

#add text for genes:
circos.text(v34a$POS[1], 0.5, v34a$GENE[1], sector.index = "chr", track.index = 1)
circos.text(v34a$POS[10], 0.5, v34a$GENE[10], sector.index = "chr", track.index = 1)
circos.text(v34a$POS[14], 0.5, v34a$GENE[14], sector.index = "chr", track.index = 1)
circos.text(v34a$POS[2], 0.5, 'Gp2', sector.index = "chr", track.index = 1)
circos.text(v34a$POS[4], 0.5, 'Gp2', sector.index = "chr", track.index = 1)
circos.text(v34b$POS[1], 0.5, v34b$GENE[1], sector.index = "chr", track.index = 1)
circos.text(v34b$POS[2], 0.5, v34b$GENE[2], sector.index = "chr", track.index = 1)

#add genome A
circos.track('chr', ylim = c(0,1),panel.fun = function(x, y) {},
             bg.col="slateblue1", bg.border=F, track.height=0.05)
circos.trackPoints('chr',-5000,0.5, col = "slateblue1",pch=15, cex = 1.4)
#coding: circle filled (16), insert/delete: square (0 0r 15 (coding)), non-coding = circle, outline (1)
for (i in 1:dim(v34a)[1]){
  circos.trackPoints('chr', v34a$POS[i], 0.5, col = 'black', pch = v34a$ptype[i], cex = 1.1)
}

#add genome B
circos.track('chr', ylim = c(0,1),panel.fun = function(x, y) {},
             bg.col="slateblue1", bg.border=F, track.height=0.05)
circos.trackPoints('chr',-5000,0.5, col = "slateblue1",pch=15, cex = 1.4)
for (i in 1:dim(v34b)[1]){
  circos.trackPoints('chr', v34b$POS[i], 0.5, col = 'black', pch = v34b$ptype[i], cex = 1.1)
}

dev.off()


#######################
#Plots for strain 39:
#read files:
v39a <- read.delim('SNP_data/39A.tsv')
v39a <- v39a[which(v39a$CHROM == "contig_1"),]
v39b <- read.delim('SNP_data/39B.tsv')
v39b <- v39b[which(v39b$CHROM == "contig_1"),]
v39c <- read.delim('SNP_data/39C.tsv')
v39c <- v39c[which(v39c$CHROM == "contig_1"),]
v39d <- read.delim('SNP_data/39D.tsv')
v39d <- v39d[which(v39d$CHROM == "contig_1"),]

png("Figures/circle39.png", width = 1500, height = 1500, units = "px", bg = "white",  res = 300)

circos.clear()
circos.par("track.height" = 0.05)
col_text <- "white"
circos.par(start.degree = 90)
circos.initialize(sectors = 'chr', xlim = c(0,g34))
circos.track(ylim=c(0, 1), panel.fun=function(x, y) {
  chr=CELL_META$sector.index
  xlim=CELL_META$xlim
  ylim=CELL_META$ylim
  #circos.axis(labels.cex = 0.6)
}, bg.col="white", bg.border=F, track.height=0.1)

#add text for genes:
circos.text(v39a$POS[1], 0.5, v39a$GENE[1], sector.index = "chr", track.index = 1)
circos.text(v39a$POS[3], 0.5, v39a$GENE[3], sector.index = "chr", track.index = 1)
circos.text(v39a$POS[4], 0.5, v39a$GENE[4], sector.index = "chr", track.index = 1)
circos.text(v39b$POS[2], 0.5, v39b$GENE[2], sector.index = "chr", track.index = 1)
circos.text(v39c$POS[1], 0.5, v39c$GENE[1], sector.index = "chr", track.index = 1)
circos.text(v39d$POS[2], 0.5, v39d$GENE[2], sector.index = "chr", track.index = 1)

#add genome A
circos.track('chr', ylim = c(0,1),panel.fun = function(x, y) {},
             bg.col="palegreen3", bg.border=F, track.height=0.05)
circos.trackPoints('chr',-5000,0.5, col = "palegreen3",pch=15, cex = 1.4)
for (i in 1:dim(v39a)[1]){
  circos.trackPoints('chr', v39a$POS[i], 0.5, col = 'black', pch = v39a$ptype[i], cex = 1.1)
}

#add genome B
circos.track('chr', ylim = c(0,1),panel.fun = function(x, y) {},
             bg.col="palegreen3", bg.border=F, track.height=0.05)
circos.trackPoints('chr',-5000,0.5, col = "palegreen3",pch=15, cex = 1.4)
for (i in 1:dim(v39b)[1]){
  circos.trackPoints('chr', v39b$POS[i], 0.5, col = 'black', pch = v39b$ptype[i], cex = 1.1)
}

#add genome C
circos.track('chr', ylim = c(0,1),panel.fun = function(x, y) {},
             bg.col="palegreen3", bg.border=F, track.height=0.05)
circos.trackPoints('chr',-5000,0.5, col = "palegreen3",pch=15, cex = 1.4)
for (i in 1:dim(v39c)[1]){
  circos.trackPoints('chr', v39c$POS[i], 0.5, col = 'black', pch = v39c$ptype[i], cex = 1.1)
}

#add genome D
circos.track('chr', ylim = c(0,1),panel.fun = function(x, y) {},
             bg.col="palegreen3", bg.border=F, track.height=0.05)
circos.trackPoints('chr',-5000,0.5, col = "palegreen3",pch=15, cex = 1.4)
for (i in 1:dim(v39d)[1]){
  circos.trackPoints('chr', v39d$POS[i], 0.5, col = 'black', pch = v39d$ptype[i], cex = 1.1)
}

dev.off()

#######################
#Plots for strain 59:
#read files:
v59a <- read.delim("SNP_data/59A.tsv")
v59a <- v59a[which(v59a$CHROM == "contig_1"),]
v59b <- read.delim("SNP_data/59B.tsv")
v59b <- v59b[which(v59b$CHROM == "contig_1"),]
v59c <- read.delim("SNP_data/59C.tsv")
v59c <- v59c[which(v59c$CHROM == "contig_1"),]
v59d <- read.delim("SNP_data/59D.tsv")
v59d <- v59d[which(v59d$CHROM == "contig_1"),]

png("Figures/circle59.png", width = 1500, height = 1500, units = "px", bg = "white",  res = 300)

circos.clear()
circos.par("track.height" = 0.05)
col_text <- "white"
circos.par(start.degree = 90)
circos.initialize(sectors = 'chr', xlim = c(0,g34))
circos.track(ylim=c(0, 1), panel.fun=function(x, y) {
  chr=CELL_META$sector.index
  xlim=CELL_META$xlim
  ylim=CELL_META$ylim
  #circos.axis(labels.cex = 0.6)
}, bg.col="white", bg.border=F, track.height=0.1)

#add text for genes:
circos.text(v59a$POS[1], 0.5, v59a$GENE[1], sector.index = "chr", track.index = 1)
circos.text(v59a$POS[2], 0.5, v59a$GENE[2], sector.index = "chr", track.index = 1)
circos.text(v59a$POS[3], 0.5, v59a$GENE[3], sector.index = "chr", track.index = 1)
circos.text(v59a$POS[5], 0.5, v59a$GENE[5], sector.index = "chr", track.index = 1)

#add genome A
circos.track('chr', ylim = c(0,1),panel.fun = function(x, y) {},
             bg.col="gold2", bg.border=F, track.height=0.05)
circos.trackPoints('chr',-5000,0.5, col = "gold2",pch=15, cex = 1.4)
for (i in 1:dim(v59a)[1]){
  circos.trackPoints('chr', v59a$POS[i], 0.5, col = 'black', pch = v59a$ptype[i], cex = 1.1)
}

#add genome B
circos.track('chr', ylim = c(0,1),panel.fun = function(x, y) {},
             bg.col="gold2", bg.border=F, track.height=0.05)
circos.trackPoints('chr',-5000,0.5, col = "gold2",pch=15, cex = 1.4)
for (i in 1:dim(v59b)[1]){
  circos.trackPoints('chr', v59b$POS[i], 0.5, col = 'black', pch = v59b$ptype[i], cex = 1.1)
}

#add genome C
circos.track('chr', ylim = c(0,1),panel.fun = function(x, y) {},
             bg.col="gold2", bg.border=F, track.height=0.05)
circos.trackPoints('chr',-5000,0.5, col = "gold2",pch=15, cex = 1.4)
for (i in 1:dim(v59c)[1]){
  circos.trackPoints('chr', v59c$POS[i], 0.5, col = 'black', pch = v59c$ptype[i], cex = 1.1)
}

#add genome D
circos.track('chr', ylim = c(0,1),panel.fun = function(x, y) {},
             bg.col="gold2", bg.border=F, track.height=0.05)
circos.trackPoints('chr',-5000,0.5, col = "gold2",pch=15, cex = 1.4)
for (i in 1:dim(v59d)[1]){
  circos.trackPoints('chr', v59d$POS[i], 0.5, col = 'black', pch = v59d$ptype[i], cex = 1.1)
}

dev.off()
```
