## Supplementary material for "The effect of antibiotic selection on collateral effects and evolvability of uropathogenic *Escherichia coli*": Code and commands: PlotsEvolvabilityManuscript.R

```
rm(list=ls(all=TRUE))
dev.off()
setwd(" ") #Add path to data files. Figures will be saved to Figures folder
library(ggplot2)
library(ggalt)
library(GGally)
library(ggpubr)
library(reshape2)
library(tidyverse)
library(ggbeeswarm)
library(growthcurver)

#EUCAST breakpoints (for oral, uncomplicated UTI):
bkp_trim <- 4
bkp_nitro <- 64  
bkp_fos <- 8 

########################################################
######### Plots for MICs for ancestors and trimR strains
########################################################

#read in MIC data
mic <- read.csv('MIC_all_tidy.csv')
mtrim <- mic[which(mic$Antibiotic=="TRIM"),]
mnitro <- mic[which(mic$Antibiotic =="NITRO"),]
mfos <- mic[which(mic$Antibiotic =="FOS"),]

#Graph for individual MICs for TRIM, ancestor and trimR mutants
level_order <- c('34','34A','34B','39','39A','39B','39C','39D','59','59A','59B','59C','59D') 
col = c('slateblue4','slateblue','slateblue', 'darkgreen',rep('palegreen3',4),'goldenrod3',rep('gold2',4))

ggplot(mtrim, aes(x=Strain, y=MIC,color= Strain ))+
  geom_quasirandom(alpha = 1, width =.25, size = 2.5 ) +
  stat_summary(fun = "mean",geom = "crossbar", width = 0.2,colour = "gray30") +
  scale_y_continuous(trans='log2', limits = c(.25,128)) +
  labs(x="Strains", y= "Trimethoprim (\U03BCg/\U03BCl)")+
  scale_color_manual(values = col) +
  scale_fill_manual(values = col) + 
  geom_hline(yintercept= bkp_trim, linetype="dotted", color = "gray20", size=.7) +
  theme(legend.position = "none", axis.text.x=element_text(size=12),
        axis.text=element_text(size=12),strip.text = element_text(size = 14)) +
  facet_grid(. ~ Strain_background, scales = "free")

ggsave("Figures/trim_mic_separate.png", width = 12, height = 4, device='png', dpi=150)

#Graph for individual MICs for NITRO, ancestor and trinR mutants
ggplot(mnitro, aes(x=Strain, y=MIC,color= Strain ))+
  geom_quasirandom(alpha = 1, width =.25, size =2.5) +
  stat_summary(fun = "mean",geom = "crossbar", width = 0.2,colour = "gray30") +
  scale_y_continuous(trans='log2', limits = c(.25,128)) +
  labs(x="strains", y= "Nitrofurantoin (\U03BCg/\U03BCl)")+
  scale_color_manual(values = col) +
  scale_fill_manual(values = col) + 
  geom_hline(yintercept= bkp_nitro, linetype="dotted", color = "gray20", size=.7) +
  theme(legend.position = "none", axis.text.x=element_text(size=12),
        axis.text=element_text(size=12),strip.text = element_text(size = 14)) +
  facet_grid(. ~ Strain_background, scales = "free")

ggsave("Figures/nitro_mic_separate.png", width = 12, height = 4, device='png', dpi=150)

#Graph for individual MICs for FOS, ancestor and trimR mutants
ggplot(mfos, aes(x=Strain, y=MIC,color= Strain ))+
  geom_quasirandom(alpha = 1, width =.25, size =2.5) +
  stat_summary(fun = "mean", geom = "crossbar", width = 0.2,colour = "gray30") +
  scale_y_continuous(trans='log2',limits = c(.25,128)) +
  labs(x="strains", y= "Fosfomycin (\U03BCg/\U03BCl)")+
  scale_color_manual(values = col) +
  scale_fill_manual(values = col) + 
  geom_hline(yintercept= bkp_fos, linetype="dotted", color = "gray20", size=.7) +
  theme(legend.position = "none", axis.text.x=element_text(size=12),
        axis.text=element_text(size=12),strip.text = element_text(size = 14)) +
  facet_grid(. ~ Strain_background, scales = "free")

ggsave("Figures/fos_mic_separate.png", width = 12, height = 4, device='png', dpi=150)


#individual MIC for TRIM, grouped by Ancestor and trimR, coloured by strain:
eb11t <- mean(mtrim[which(mtrim$Ancestor == "Ancestor"),5])-sd(mtrim[which(mtrim$Ancestor == "Ancestor"),5])/sqrt(length(which(mtrim$Ancestor=="Ancestor")))
eb12t <- mean(mtrim[which(mtrim$Ancestor == "Ancestor"),5])+sd(mtrim[which(mtrim$Ancestor == "Ancestor"),5])/sqrt(length(which(mtrim$Ancestor=="Ancestor")))
eb21t <- mean(mtrim[which(mtrim$Ancestor == "Trim R"),5])-sd(mtrim[which(mtrim$Ancestor == "Trim R"),5])/sqrt(length(which(mtrim$Ancestor=="Trim R")))
eb22t <- mean(mtrim[which(mtrim$Ancestor == "Trim R"),5])+sd(mtrim[which(mtrim$Ancestor == "Trim R"),5])/sqrt(length(which(mtrim$Ancestor=="Trim R")))
p1<-ggplot(mtrim, aes(x=Ancestor, y=MIC,color= Strain ))+
  geom_quasirandom(alpha = 1, width =.25, size =2.5) +
  geom_segment(aes(x = 1, y = eb11t, xend = 1,yend = eb12t) ,colour = "gray30") +
  geom_segment(aes(x = 2, y = eb21t, xend = 2,yend = eb22t) ,colour = "gray30") +
  geom_segment(aes(x = 0.9, y = mean(mtrim[which(mtrim$Ancestor == "Ancestor"),5]), xend = 1.1 ,yend = mean(mtrim[which(mtrim$Ancestor == "Ancestor"),5])) ,colour = "gray30",size=1.5) +
  geom_segment(aes(x = 1.9, y = mean(mtrim[which(mtrim$Ancestor == "Trim R"),5]), xend = 2.1, yend = mean(mtrim[which(mtrim$Ancestor == "Trim R"),5])) ,colour = "gray30",size=1.5) +
  scale_y_continuous(trans='log2', limits = c(.25,128)) +
  labs( y= "Trimethoprim (\U03BCg/\U03BCl)",x='')+
  scale_color_manual(values = col) + 
  scale_fill_manual(values = col) + 
  geom_hline(yintercept= bkp_trim, linetype="dotted", color = "gray20", size=.7) +
  theme(legend.position = "none", axis.text.x=element_text(size=12),
        axis.text=element_text(size=12))

#individual MIC for NITRO, grouped by Ancestor and trimR, coloured by strain:
eb11n <- mean(mnitro[which(mnitro$Ancestor == "Ancestor"),5])-sd(mnitro[which(mnitro$Ancestor == "Ancestor"),5])/sqrt(length(which(mnitro$Ancestor=="Ancestor")))
eb12n <- mean(mnitro[which(mnitro$Ancestor == "Ancestor"),5])+sd(mnitro[which(mnitro$Ancestor == "Ancestor"),5])/sqrt(length(which(mnitro$Ancestor=="Ancestor")))
eb21n <- mean(mnitro[which(mnitro$Ancestor == "Trim R"),5])-sd(mnitro[which(mnitro$Ancestor == "Trim R"),5])/sqrt(length(which(mnitro$Ancestor=="Trim R")))
eb22n <- mean(mnitro[which(mnitro$Ancestor == "Trim R"),5])+sd(mnitro[which(mnitro$Ancestor == "Trim R"),5])/sqrt(length(which(mnitro$Ancestor=="Trim R")))
p2<-ggplot(mnitro, aes(x=Ancestor, y=MIC,color= Strain ))+
  geom_quasirandom(alpha = 1, width =.25, size =2.5) +
  geom_segment(aes(x = 1, y = eb11n, xend = 1,yend = eb12n) ,colour = "gray30") +
  geom_segment(aes(x = 2, y = eb21n, xend = 2,yend = eb22n) ,colour = "gray30") +
  geom_segment(aes(x = 0.9, y = mean(mnitro[which(mnitro$Ancestor == "Ancestor"),5]), xend = 1.1 ,yend = mean(mnitro[which(mnitro$Ancestor == "Ancestor"),5])) ,colour = "gray30",size=1.5) +
  geom_segment(aes(x = 1.9, y = mean(mnitro[which(mnitro$Ancestor == "Trim R"),5]), xend = 2.1, yend = mean(mnitro[which(mnitro$Ancestor == "Trim R"),5])) ,colour = "gray30",size=1.5) +
  scale_y_continuous(trans='log2', limits = c(.25,128)) +
  labs(y = "Nitrofurantoin (\U03BCg/\U03BCl)",x="")+
  scale_color_manual(values = col) +
  scale_fill_manual(values = col) + 
  geom_hline(yintercept= bkp_nitro, linetype="dotted", color = "gray20", size=.7) +
  theme(legend.position = "none", axis.text.x=element_text(size=12),
        axis.text=element_text(size=12))

#individual MIC for FOS, grouped by Ancestor and trimR, coloured by strain:
eb11f <- mean(mfos[which(mfos$Ancestor == "Ancestor"),5])-sd(mfos[which(mfos$Ancestor == "Ancestor"),5])/sqrt(length(which(mfos$Ancestor=="Ancestor")))
eb12f <- mean(mfos[which(mfos$Ancestor == "Ancestor"),5])+sd(mfos[which(mfos$Ancestor == "Ancestor"),5])/sqrt(length(which(mfos$Ancestor=="Ancestor")))
eb21f <- mean(mfos[which(mfos$Ancestor == "Trim R"),5])-sd(mfos[which(mfos$Ancestor == "Trim R"),5])/sqrt(length(which(mfos$Ancestor=="Trim R")))
eb22f <- mean(mfos[which(mfos$Ancestor == "Trim R"),5])+sd(mfos[which(mfos$Ancestor == "Trim R"),5])/sqrt(length(which(mfos$Ancestor=="Trim R")))
p3<-ggplot(mfos, aes(x=Ancestor, y=MIC,color= Strain ))+#, fill = group, color =group)) +
  geom_quasirandom(alpha = 1, width =.25, size =2.5) +
  geom_segment(aes(x = 1, y = eb11f, xend = 1,yend = eb12f) ,colour = "gray30") +
  geom_segment(aes(x = 2, y = eb21f, xend = 2,yend = eb22f) ,colour = "gray30") +
  geom_segment(aes(x = 0.9, y = mean(mfos[which(mfos$Ancestor == "Ancestor"),5]), xend = 1.1 ,yend = mean(mfos[which(mfos$Ancestor == "Ancestor"),5])) ,colour = "gray30",size=1.5) +
  geom_segment(aes(x = 1.9, y = mean(mfos[which(mfos$Ancestor == "Trim R"),5]), xend = 2.1, yend = mean(mfos[which(mfos$Ancestor == "Trim R"),5])) ,colour = "gray30",size=1.5) +
  scale_y_continuous(trans='log2', limits = c(.25,128)) +
  labs(y= "Fosfomycin (\U03BCg/\U03BCl)",x="")+
  scale_color_manual(values = col) +
  scale_fill_manual(values = col) +  
  geom_hline(yintercept= bkp_fos, linetype="dotted", color = "gray20", size=.7) +
  theme(legend.position = "none", axis.text.x=element_text(size=12),
        axis.text=element_text(size=12))

ggarrange(p1,p2,p3,nrow =1)
ggsave("Figures/MIC_anc&ev.png", width = 12, height = 4, device='png', dpi=150)

##############################
############### MSW graphs
##############################

#read data
res <- read.csv('MSW_all_tidy.csv')
#make sure data is numeric and separate
res$MIC <- as.numeric(res$MIC)
res$MBC <- as.numeric(res$MBC)
res$MPC <- as.numeric(res$MPC)
rtrim <- res[which(res$Antibiotic=="TRIM"),]
rnitro <- res[which(res$Antibiotic =="NITRO"),]
rfos <- res[which(res$Antibiotic =="FOS"),]


#MSW  overall graph grouped in ancestors and trim-resistant for trim, nitro and fos.
#calculatate averages for ancestor and trimR:
trim_ac <- colMeans(rtrim[which(rtrim$Ancestor==1),5:7],na.rm = TRUE)
trim_ev <- colMeans(rtrim[which(rtrim$Ancestor==0),5:7],na.rm = TRUE)

nitro_ac <- colMeans(rnitro[which(rnitro$Ancestor==1),5:7],na.rm = TRUE)
nitro_ev <- colMeans(rnitro[which(rnitro$Ancestor==0),5:7],na.rm = TRUE)

fos_ac <- colMeans(rfos[which(rfos$Ancestor==1),5:7],na.rm = TRUE)
fos_ev <- colMeans(rfos[which(rfos$Ancestor==0),5:7],na.rm = TRUE)

tdft <- as.data.frame(rbind(trim_ev,trim_ac))
tdft <- cbind(c("Trim R", "Ancestor"),tdft)
colnames(tdft) <- c('strain',colnames(res)[5:7])

#TRIM. MPC
p1 <- ggplot(tdft, aes(y = tdft$strain, x = tdft$MIC , xend = tdft$MPC)) + 
  geom_dumbbell(size=3, color="#9e9e9e",colour_x = "#95AAD3", colour_xend = "darkblue", dot_guide=FALSE) +
  labs(x= expression( "Trimethoprim (\U03BCg/\U03BCl)")) +
  scale_x_continuous(trans='log2', limits = c(.25,256)) +
  geom_vline(xintercept= bkp_trim, linetype="dotted", color = "gray20", size=.7) +
  theme(panel.grid.minor = element_blank(),
        axis.title.x = element_text(size = 16),
        axis.text.x = element_text(size = 14),
        axis.title.y = element_blank(),
        axis.text.y = element_text(size=16),
        axis.line = element_line(linewidth = .5, colour = "grey80"))

#NITRO
tdfn <- as.data.frame(rbind(nitro_ev,nitro_ac))
tdfn <- cbind(c("Trim R", "Ancestor"),tdfn)
colnames(tdfn) <- c('strain',colnames(res)[5:7])

#NITRO. MPC
p2<-ggplot(tdfn, aes(y = tdfn$strain, x = tdfn$MIC , xend = tdfn$MPC)) + 
  geom_dumbbell(size=3, color="#9e9e9e",colour_x = "#F6BF93", colour_xend = "orangered3", dot_guide=FALSE) +
  labs(x=  "Nitrofurantoin (\U03BCg/\U03BCl)") +
  scale_x_continuous(trans='log2', limits = c(.25,128)) +
  geom_vline(xintercept= bkp_nitro, linetype="dotted", color = "gray20", size=.7) +
  theme(panel.grid.minor = element_blank(),
        axis.title.x = element_text(size = 16),
        axis.text.x = element_text(size = 14),
        axis.title.y = element_blank(),
        axis.text.y = element_text(size=16),
        axis.line = element_line(linewidth = .5, colour = "grey80"))

#FOS
tdff <- as.data.frame(rbind(fos_ev,fos_ac))
tdff <- cbind(c("Trim R", "Ancestor"),tdff)
colnames(tdff) <- c('strain',colnames(res)[5:7])

p3<-ggplot(tdff, aes(y = tdff$strain, x = tdff$MIC , xend = tdff$MPC)) + 
  geom_dumbbell(size=3, color="#9e9e9e",colour_x = "palegreen", colour_xend = "darkgreen", dot_guide=FALSE) +
  labs(x=  "Fosfomycin (\U03BCg/\U03BCl)") +
  scale_x_continuous(trans='log2', limits = c(.25,128)) +
  geom_vline(xintercept= bkp_fos, linetype="dotted", color = "gray20", size=.7) +
  theme(panel.grid.minor = element_blank(),
        axis.title.x = element_text(size = 16),
        axis.text.x = element_text(size = 14),
        axis.title.y = element_blank(),
        axis.text.y = element_text(size=16),
        axis.line = element_line(linewidth = .5, colour = "grey80"))

ggarrange(p1,p2,p3, nrow=1)
ggsave("Figures/MSW_anc&ev.png", width = 12, height = 4, device='png', dpi=150)

# MSW graph of individual MSW, with individual replicates

#TRIM. MPC individual replicates
ggplot(rtrim, aes(y = Position, x = MIC , xend = MPC)) + 
  geom_dumbbell(size=3, color="#9e9e9e",colour_x = "#95AAD3", colour_xend = "darkblue", dot_guide=FALSE) +
  labs(x= "Trimethoprim (\U03BCg/\U03BCl)",
       y = "     Anc            A            B            C            D     ") +
  scale_x_continuous(trans='log2', limits = c(.25,512)) +
  geom_vline(xintercept= bkp_trim, linetype="dotted", color = "gray20", size=.7) +
  theme(panel.grid.minor = element_blank(),
        axis.title.x = element_text(size = 14),
        axis.text.x = element_text(size = 14),
        axis.title.y = element_text(size = 14),
        axis.text.y = element_blank(),
        strip.text = element_text(size = 14),
        axis.line = element_line(linewidth = .5, colour = "grey80")) +
  facet_grid(. ~ Strain_background)
ggsave("Figures/trim_mpc.png", width = 12, height = 5, device='png', dpi=150)

#NITRO MPC individual replicates
ggplot(rnitro, aes(y = Position, x = MIC , xend = MPC)) + 
  geom_dumbbell(size=3, color="#9e9e9e",colour_x = "#F6BF93", colour_xend = "orangered3", dot_guide=FALSE) +
  labs(x= "Nitrofurantoin (\U03BCg/\U03BCl)",
       y = "     Anc            A            B            C            D     ") +
  scale_x_continuous(trans='log2', limits = c(.25,128)) +
  geom_vline(xintercept= bkp_nitro, linetype="dotted", color = "gray20", size=.7) +
  theme(panel.grid.minor = element_blank(),
        axis.title.x = element_text(size = 14),
        axis.text.x = element_text(size = 14),
        axis.title.y = element_text(size = 14),
        axis.text.y = element_blank(),
        strip.text = element_text(size = 14),
        axis.line = element_line(linewidth = .5, colour = "grey80")) +
  facet_grid(. ~ Strain_background)
ggsave("Figures/nitro_mpc.png", width = 12, height = 5, device='png', dpi=150)

#FOS MPC individual replicates
ggplot(rfos, aes(y = Position, x = MIC , xend = MPC)) + 
  geom_dumbbell(size=3, color="#9e9e9e",colour_x = "palegreen", colour_xend = "darkgreen", dot_guide=FALSE) +
  labs(x= "Fosfomycin (\U03BCg/\U03BCl)",
       y = "     Anc            A            B            C            D     ") +
  scale_x_continuous(trans='log2', limits = c(.25,128)) +
  geom_vline(xintercept= bkp_fos, linetype="dotted", color = "gray20", size=.7) +
  theme(panel.grid.minor = element_blank(),
        axis.title.x = element_text(size = 14),
        axis.text.x = element_text(size = 14),
        axis.title.y = element_text(size = 14),
        axis.text.y = element_blank(),
        strip.text = element_text(size = 14),
        axis.line = element_line(linewidth = .5, colour = "grey80")) +
  facet_grid(. ~ Strain_background)
ggsave("Figures/fos_mpc.png", width = 12, height = 5, device='png', dpi=150)

######################################
################# Fitness plots
######################################

#read OD data:
od1 <- read.csv('ODplate1.csv')
od2 <- read.csv('ODplate2.csv')
od3 <- read.csv('ODplate3.csv')

#Get growth curve parameters (gowthcurver package, will also output pdf of fits)
gc1 <- SummarizeGrowthByPlate(od1,plot_fit = TRUE, plot_file = "Figures/gcfit1.pdf")
gc2 <- SummarizeGrowthByPlate(od2,plot_fit = TRUE, plot_file = "Figures/gcfit2.pdf")
gc3 <- SummarizeGrowthByPlate(od3,plot_fit = TRUE, plot_file = "Figures/gcfit3.pdf")

gca <- rbind(gc1,gc2,gc3)
write.csv(gca, "GrowthCurveFitData.csv") #File edited to add colums with strain information
#read edited file:
gca <- read.csv("GrowthCurveFitDataE.csv",row.names = 1)
colnames(gca)[8] <- "AUC"

#Individual plots
#generation time
ggplot(gca, aes(x=factor(Strain,level = level_order), y=t_gen,color= factor(Strain) ))+#, fill = group, color =group)) +
  geom_quasirandom(alpha = 1, width =.2) +
  stat_summary(fun = "mean",geom = "crossbar", width = 0.3,colour = "black") +
  labs(x="strains", y="generation time (min)") +
  scale_color_manual(values = col) +
  scale_fill_manual(values = col)  +
  theme(legend.position = "none", axis.text.x=element_text(size=12),
        axis.text=element_text(size=12),strip.text = element_text(size = 14)) +
  facet_grid(. ~ Strain_background, scales = "free")
ggsave("Figures/gt_separate.png", width = 12, height = 4, device='png', dpi=150)

#k
ggplot(gca, aes(x=factor(Strain,level = level_order), k,color= factor(Strain) ))+#, fill = group, color =group)) +
  geom_quasirandom(alpha = 1, width =.2) +
  stat_summary(fun = "mean",geom = "crossbar", width = 0.3,colour = "black") +
  labs(x="strains", y="k") +
  scale_color_manual(values = col) +
  scale_fill_manual(values = col)  +
  theme(legend.position = "none", axis.text.x=element_text(size=12),
        axis.text=element_text(size=12),strip.text = element_text(size = 14)) +
  facet_grid(. ~ Strain_background, scales = "free")
ggsave("Figures/k_separate.png", width = 12, height = 4, device='png', dpi=150)

#AUC
ggplot(gca, aes(x=factor(Strain,level = level_order), y=AUC,color= factor(Strain) ))+#, fill = group, color =group)) +
  geom_quasirandom(alpha = 1, width =.2) +
  stat_summary(fun = "mean",geom = "crossbar", width = 0.3,colour = "gray20") +
  labs(x="strains", y="AUC") +
  scale_color_manual(values = col) +
  scale_fill_manual(values = col)  +
  theme(legend.position = "none", axis.text.x=element_text(size=12),
        axis.text=element_text(size=12),strip.text = element_text(size = 14)) +
  facet_grid(. ~ Strain_background, scales = "free")
ggsave("Figures/auc_separate.png", width = 12, height = 4, device='png', dpi=150)

#R
ggplot(gca, aes(x=factor(Strain,level = level_order), y=r,color= factor(Strain) ))+#, fill = group, color =group)) +
  geom_quasirandom(alpha = 1, width =.2) +
  stat_summary(fun = "mean",geom = "crossbar", width = 0.3,colour = "gray20") +
  labs(x="strains", y="r") +
  scale_color_manual(values = col) +
  scale_fill_manual(values = col)  +
  theme(legend.position = "none", axis.text.x=element_text(size=12),
        axis.text=element_text(size=12),strip.text = element_text(size = 14)) +
  facet_grid(. ~ Strain_background, scales = "free")
ggsave("Figures/r_separate.png", width = 12, height = 4, device='png', dpi=150)

# relative fitness combined strains

#calculate relative fitness:
#average values for ancestors
av34 <- colSums(gca[which(gca$Strain =="34"),2:9])/6
av39 <- colSums(gca[which(gca$Strain =="39"),2:9])/6
av59 <- colSums(gca[which(gca$Strain =="59"),2:9])/6

gcr <- gca[which(gca$Ancestor == "Trim R"),]
#gcr <-gca
gcr[which(gcr$Strain_background == 34),2:9] <- t(t(gcr[which(gcr$Strain_background == 34),2:9])/av34)
gcr[which(gcr$Strain_background == 39),2:9] <- t(t(gcr[which(gcr$Strain_background == 39),2:9])/av39)
gcr[which(gcr$Strain_background == 59),2:9] <- t(t(gcr[which(gcr$Strain_background == 59),2:9])/av59)

##########
gcrp <- gcr[,c(1,2,4,8,12)] #select variables to plot (k,r, AUC) and strain,replicate ID
gcrp[,5] <- as.character(gcrp[,5]) #ID as text, not number
gcrp <- melt(gcrp,c(1,5)) #reformat table for ggplot
col2 = c('slateblue','slateblue',rep('palegreen3',4),rep('gold2',4))

ggplot(gcrp, aes(x=variable, y= value,color= Strain))+
  geom_quasirandom(alpha = 1, width =.25, size =2) +
  stat_summary(fun.data=mean_sdl, fun.args = list(mult=1), geom="errorbar", color="gray30", width=0.3) +
  stat_summary(fun.y=mean, geom="crossbar", color="gray30",width=0.2)+
  labs(y= "Relative fitness",x="")+
  scale_color_manual(values = col2) +
  scale_fill_manual(values = col2) +  
  geom_hline(yintercept= 1, linetype="dotted", color = "gray20", size=.7) +
  theme(legend.position = "none", axis.text.x=element_text(size=12),
        axis.text=element_text(size=12))
ggsave("Figures/RelativeFit_R.png", width = 6, height = 5, device='png', dpi=150)

#Version 2, ancestor and r:
gcn <- gca
gcn[which(gcn$Strain_background == 34),2:9] <- t(t(gcn[which(gcn$Strain_background == 34),2:9])/av34)
gcn[which(gcn$Strain_background == 39),2:9] <- t(t(gcn[which(gcn$Strain_background == 39),2:9])/av39)
gcn[which(gcn$Strain_background == 59),2:9] <- t(t(gcn[which(gcn$Strain_background == 59),2:9])/av59)

##########
gcrp <- gcr[,c(1,2,4,8,11,12)]
gcrp[,5] <- as.character(gcrp[,5])
gcrp <- melt(gcrp,c(1,5))

p1<-ggplot(gcn, aes(x=Ancestor, y= k,color= Strain))+
  geom_quasirandom(alpha = 1, width =.25, size =2) +
  #scale_y_continuous(trans='log2', limits = c(.25,2)) +
  stat_summary(fun.data=mean_sdl, fun.args = list(mult=1), geom="errorbar", color="gray30", width=0.3) +
  stat_summary(fun.y=mean, geom="crossbar", color="gray30",width=0.2)+
  labs(y= "Relative k",x="")+
  scale_color_manual(values = col) +
  scale_fill_manual(values = col) +  
  geom_hline(yintercept= 1, linetype="dotted", color = "gray20", size=.7) +
  theme(legend.position = "none", axis.text.x=element_text(size=12),
        axis.text=element_text(size=12))

p2<-ggplot(gcn, aes(x=Ancestor, y= r,color= Strain))+
  geom_quasirandom(alpha = 1, width =.25, size =2) +
  #scale_y_continuous(trans='log2', limits = c(.25,2)) +
  stat_summary(fun.data=mean_sdl, fun.args = list(mult=1), geom="errorbar", color="gray30", width=0.3) +
  stat_summary(fun.y=mean, geom="crossbar", color="gray30",width=0.2)+
  labs(y= "Relative r",x="")+
  scale_color_manual(values = col) +
  scale_fill_manual(values = col) +  
  geom_hline(yintercept= 1, linetype="dotted", color = "gray20", size=.7) +
  theme(legend.position = "none", axis.text.x=element_text(size=12),
        axis.text=element_text(size=12))

p3<-ggplot(gcn, aes(x=Ancestor, y= AUC,color= Strain))+
  geom_quasirandom(alpha = 1, width =.25, size =2) +
  #scale_y_continuous(trans='log2', limits = c(.25,2)) +
  stat_summary(fun.data=mean_sdl, fun.args = list(mult=1), geom="errorbar", color="gray30", width=0.3) +
  stat_summary(fun.y=mean, geom="crossbar", color="gray30",width=0.2)+
  labs(y= "Relative AUC",x="")+
  scale_color_manual(values = col) +
  scale_fill_manual(values = col) +  
  geom_hline(yintercept= 1, linetype="dotted", color = "gray20", size=.7) +
  theme(legend.position = "none", axis.text.x=element_text(size=12),
        axis.text=element_text(size=12))
  'kgn'
ggarrange(p1,p2,p3,nrow =1)
ggsave("Figures/RelativeFit_AandR.png", width = 12, height = 4, device='png', dpi=150)

##################################################### 
################## Population establishment
###################################################### 

#read in data
pep <- read.csv('PEP_tidy.csv')
pnitro <- pep[which(pep$Antibiotic =="NITRO"),]
pfos <- pep[which(pep$Antibiotic =="FOS"),]

# Plots for individual strains, NITRO
ggplot(pnitro, aes(x=Strain, y=WellsGrowth,color= Strain ))+
  geom_quasirandom(alpha = 1, width =.25, size =2.5) +
  stat_summary(fun = "mean",geom = "crossbar", width = 0.2,colour = "gray30") +
  labs(x="strains", y= "Wells with growth")+
  scale_color_manual(values = col) +
  scale_fill_manual(values = col) + 
  theme(legend.position = "none", axis.text.x=element_text(size=12),
        axis.text=element_text(size=12),strip.text = element_text(size = 14)) +
  facet_grid(. ~ Strain_background, scales = "free")

ggsave("Figures/nitro_pep_separate.png", width = 12, height = 4, device='png', dpi=150)

# Plots for individual strains, FOS
ggplot(pfos, aes(x=Strain, y=WellsGrowth,color= Strain ))+
  geom_quasirandom(alpha = 1, width =.25, size =2.5) +
  stat_summary(fun = "mean", geom = "crossbar", width = 0.2,colour = "gray30") +
  labs(x="strains", y= "Wells with growth")+
  scale_color_manual(values = col) +
  scale_fill_manual(values = col) + 
  theme(legend.position = "none", axis.text.x=element_text(size=12),
        axis.text=element_text(size=12),strip.text = element_text(size = 14)) +
  facet_grid(. ~ Strain_background, scales = "free")

ggsave("Figures/fos_pep_separate.png", width = 12, height = 4, device='png', dpi=150)

#graph of all data grouped by ancestors and trim-resistant for nitro and fos
# different colours for the different ancestors/mutants 
#NITRO
eb11n <- mean(pnitro[which(pnitro$Ancestor == "Ancestor"),5])-sd(pnitro[which(pnitro$Ancestor == "Ancestor"),5])/sqrt(length(which(pnitro$Ancestor=="Ancestor")))
eb12n <- mean(pnitro[which(pnitro$Ancestor == "Ancestor"),5])+sd(pnitro[which(pnitro$Ancestor == "Ancestor"),5])/sqrt(length(which(pnitro$Ancestor=="Ancestor")))
eb21n <- mean(pnitro[which(pnitro$Ancestor == "Trim R"),5])-sd(pnitro[which(pnitro$Ancestor == "Trim R"),5])/sqrt(length(which(pnitro$Ancestor=="Trim R")))
eb22n <- mean(pnitro[which(pnitro$Ancestor == "Trim R"),5])+sd(pnitro[which(pnitro$Ancestor == "Trim R"),5])/sqrt(length(which(pnitro$Ancestor=="Trim R")))
p1<-ggplot(pnitro, aes(x=Ancestor, y=WellsGrowth,color= Strain ))+#, fill = group, color =group)) +
  geom_quasirandom(alpha = 1, width =.25, size =2.5) +
  geom_segment(aes(x = 1, y = eb11n, xend = 1,yend = eb12n) ,colour = "gray30") +
  geom_segment(aes(x = 2, y = eb21n, xend = 2,yend = eb22n) ,colour = "gray30") +
  geom_segment(aes(x = 0.9, y = mean(pnitro[which(pnitro$Ancestor == "Ancestor"),5]), xend = 1.1 ,yend = mean(pnitro[which(pnitro$Ancestor == "Ancestor"),5])) ,colour = "gray30",size=1.5) +
  geom_segment(aes(x = 1.9, y = mean(pnitro[which(pnitro$Ancestor == "Trim R"),5]), xend = 2.1, yend = mean(pnitro[which(pnitro$Ancestor == "Trim R"),5])) ,colour = "gray30",size=1.5) +
  labs(y = "Wells with growth",x="")+
  scale_color_manual(values = col) +
  scale_fill_manual(values = col) + 
  theme(legend.position = "none", axis.text.x=element_text(size=12),
        axis.text=element_text(size=12))

#FOS
eb11f <- mean(pfos[which(pfos$Ancestor == "Ancestor"),5])-sd(pfos[which(pfos$Ancestor == "Ancestor"),5])/sqrt(length(which(pfos$Ancestor=="Ancestor")))
eb12f <- mean(pfos[which(pfos$Ancestor == "Ancestor"),5])+sd(pfos[which(pfos$Ancestor == "Ancestor"),5])/sqrt(length(which(pfos$Ancestor=="Ancestor")))
eb21f <- mean(pfos[which(pfos$Ancestor == "Trim R"),5])-sd(pfos[which(pfos$Ancestor == "Trim R"),5])/sqrt(length(which(pfos$Ancestor=="Trim R")))
eb22f <- mean(pfos[which(pfos$Ancestor == "Trim R"),5])+sd(pfos[which(pfos$Ancestor == "Trim R"),5])/sqrt(length(which(pfos$Ancestor=="Trim R")))
p2<-ggplot(pfos, aes(x=Ancestor, y=WellsGrowth,color= Strain ))+#, fill = group, color =group)) +
  geom_quasirandom(alpha = 1, width =.25, size =2.5) +
  geom_segment(aes(x = 1, y = eb11f, xend = 1,yend = eb12f) ,colour = "gray30") +
  geom_segment(aes(x = 2, y = eb21f, xend = 2,yend = eb22f) ,colour = "gray30") +
  geom_segment(aes(x = 0.9, y = mean(pfos[which(pfos$Ancestor == "Ancestor"),5]), xend = 1.1 ,yend = mean(pfos[which(pfos$Ancestor == "Ancestor"),5])) ,colour = "gray30",size=1.5) +
  geom_segment(aes(x = 1.9, y = mean(pfos[which(pfos$Ancestor == "Trim R"),5]), xend = 2.1, yend = mean(pfos[which(pfos$Ancestor == "Trim R"),5])) ,colour = "gray30",size=1.5) +
  labs(y= "Wells with growth",x="")+
  scale_color_manual(values = col) +
  scale_fill_manual(values = col) +  
  theme(legend.position = "none", axis.text.x=element_text(size=12),
        axis.text=element_text(size=12))

ggarrange(p1,p2,nrow =1)
ggsave("Figures/PEP_anc&ev.png", width = 8, height = 4, device='png', dpi=150)


#########################################################################
######## Statistical tests: 
######## Figure 1
F1A: (trim)
kruskal.test(MIC ~ Ancestor, data =mtrim)
# Kruskal-Wallis rank sum test
# 
# data:  MIC by Ancestor
# Kruskal-Wallis chi-squared = 20.336, df = 1, p-value = 6.498e-06

mean(mtrim[mtrim$Ancestor == "Ancestor", "MIC"])
mean(mtrim[mtrim$Ancestor == "Trim R", "MIC"])
F1B: nitro
kruskal.test(MIC ~ Ancestor, data =mnitro)
# Kruskal-Wallis rank sum test
#   
# data:  MIC by Ancestor
# Kruskal-Wallis chi-squared = 21.446, df = 1, p-value = 3.639e-06

F1C
kruskal.test(MIC ~ Ancestor, data =mfos)
# Kruskal-Wallis rank sum test
# 
# data:  MIC by Ancestor
# Kruskal-Wallis chi-squared = 0.46897, df = 1, p-value = 0.4935

# #Statistics for MSW: Ancestor vs evolved for each antibiotic (FIg2)
#MIC
kruskal.test(MIC ~ Ancestor, data =rtrim)
# Kruskal-Wallis rank sum test
# 
# data:  MIC by Ancestor
# Kruskal-Wallis chi-squared = 20.336, df = 1, p-value = 6.498e-06

kruskal.test(MIC ~ Ancestor, data =rnitro)
# Kruskal-Wallis rank sum test
# 
# data:  MIC by Ancestor
# Kruskal-Wallis chi-squared = 21.446, df = 1, p-value = 3.639e-06
mean(rnitro[rnitro$Ancestor == 0, "MIC"],na.rm = TRUE) #28.8
mean(rnitro[rnitro$Ancestor == 1, "MIC"],na.rm = TRUE) #6.666667

kruskal.test(MIC ~ Ancestor, data =rfos)
# Kruskal-Wallis rank sum test
# 
# data:  MIC by Ancestor
# Kruskal-Wallis chi-squared = 0.46897, df = 1, p-value = 0.4935
mean(rfos[rfos$Ancestor == 1, "MIC"])
mean(rfos[rfos$Ancestor == 0, "MIC"],na.rm = TRUE)

######## Figure 2
#MPC
kruskal.test(MPC ~ Ancestor, data =rtrim)
# Kruskal-Wallis rank sum test
# 
# data:  MPC by Ancestor
# Kruskal-Wallis chi-squared = 20.609, df = 1, p-value = 5.634e-06
mean(rtrim[rtrim$Ancestor == 1, "MPC"])

kruskal.test(MPC ~ Ancestor, data =rnitro)
# Kruskal-Wallis rank sum test
# 
# data:  MPC by Ancestor
# Kruskal-Wallis chi-squared = 7.6235, df = 1, p-value = 0.005761
kruskal.test(MPC ~ Ancestor, data =rfos)
# Kruskal-Wallis rank sum test
# 
# data:  MPC by Ancestor
# Kruskal-Wallis chi-squared = 0.74126, df = 1, p-value = 0.3893


#
kruskal.test((MPC-MIC) ~ Ancestor, data =rtrim)
# Kruskal-Wallis rank sum test
# 
# data:  (MPC - MIC) by Ancestor
# Kruskal-Wallis chi-squared = 20.33, df = 1, p-value = 6.516e-06
kruskal.test((MPC-MIC) ~ Ancestor, data =rnitro)
# Kruskal-Wallis rank sum test
# 
# data:  (MPC - MIC) by Ancestor
# Kruskal-Wallis chi-squared = 19.825, df = 1, p-value = 8.485e-06
kruskal.test((MPC-MIC) ~ Ancestor, data =rfos)
# Kruskal-Wallis rank sum test
# 
# data:  (MPC - MIC) by Ancestor
# Kruskal-Wallis chi-squared = 0.001287, df = 1, p-value = 0.9714


######## Figure 4
###k
#test for normality:
shapiro.test(as.numeric(gcrp[which(gcrp$variable == 'k'),4]))
# Shapiro-Wilk normality test
# 
# data:  as.numeric(gcrp[which(gcrp$variable == "k"), 4])
# W = 0.97525, p-value = 0.2612

#Normal, so t test: 
t.test(as.numeric(gcrp[which(gcrp$variable == 'k'),4])-1)
# data:  as.numeric(gcrp[which(gcrp$variable == "k"), 4]) - 1
# t = -5.8202, df = 59, p-value = 2.566e-07
# alternative hypothesis: true mean is not equal to 0
# 95 percent confidence interval:
#   -0.09326406 -0.04554201
# sample estimates:
#   mean of x 
# -0.06940304 

mean(as.numeric(gcrp[gcrp$variable == "r", "value"]))

###r
shapiro.test(as.numeric(gcrp[which(gcrp$variable == 'r'),4]))
# Shapiro-Wilk normality test
# 
# data:  as.numeric(gcrp[which(gcrp$variable == "r"), 4])
# W = 0.91582, p-value = 0.0005226

#Not normal, so Wilcoxon:
wilcox.test(as.numeric(gcrp[which(gcrp$variable == 'r'),4])-1)
# Wilcoxon signed rank test with continuity correction
# 
# data:  as.numeric(gcrp[which(gcrp$variable == "r"), 4]) - 1
# V = 886, p-value = 0.8338
# alternative hypothesis: true location is not equal to 0


shapiro.test(as.numeric(gcrp[which(gcrp$variable == 'AUC'),4]))
# Shapiro-Wilk normality test
# 
# data:  as.numeric(gcrp[which(gcrp$variable == "AUC"), 4])
# W = 0.98576, p-value = 0.7098

#Normal, so t test:
t.test(as.numeric(gcrp[which(gcrp$variable == 'AUC'),4])-1)
# data:  as.numeric(gcrp[which(gcrp$variable == "AUC"), 4]) - 1
# t = -7.3679, df = 59, p-value = 6.44e-10
# alternative hypothesis: true mean is not equal to 0
# 95 percent confidence interval:
#   -0.12360428 -0.07080605
# sample estimates:
#   mean of x 
# -0.09720516 

#Population establishment

#nitro

kruskal.test(WellsGrowth ~ Ancestor, data =pnitro)

# Kruskal-Wallis rank sum test
# 
# data:  WellsGrowth by Ancestor
# Kruskal-Wallis chi-squared = 0.18837, df = 1, p-value = 0.6643


kruskal.test(WellsGrowth ~ Ancestor, data =pfos)

# Kruskal-Wallis rank sum test
# 
# data:  WellsGrowth by Ancestor
# Kruskal-Wallis chi-squared = 0.0044544, df = 1, p-value = 0.9468
```
